# BATF3 controls the homeostatic maintenance and function of innate-like γδ T cells

**DOI:** 10.64898/2025.12.16.694583

**Authors:** Pedro H. Papotto, Elena Hernández-Garcia, Ángela Zarco-Cuadrillero, Ana Redondo-Urzainqui, James E. Parkinson, Rúben G.R. Pinheiro, Andrew S. MacDonald, David Sancho, Bruno Silva-Santos, Judith E. Allen, Adrian C. Hayday, Salvador Iborra, Miguel Muñoz-Ruiz

## Abstract

γδ T cells compose an evolutionarily conserved lineage of lymphocytes, with both adaptive- and innate-like characteristics, contributing to tissue homeostasis, immune surveillance, and rapid responses to stress and infection. While their functional diversity and tissue-specific roles are tightly regulated by transcriptional networks, the underlying molecular mechanisms remain incompletely understood. The transcription factor basic leucine zipper ATF-like transcription factor 3 (BATF3) plays a central role in the development of conventional type 1 dendritic cells (cDC1s). Here, we unveil BATF3 as a critical cell-intrinsic regulator of the homeostasis, functional specialization, and tissue distribution of γδ T cells. *Batf3*-deficient mice display an altered composition of γδ T cell subsets, with a marked decrease in the numbers of innate-like γδ T cells across multiple organs when compared to their wild-type counterparts, independently of cDC1s. Loss of BATF3 impacts not only cell survival but also IL-17 production after γδ T cells complete their thymic development. Mechanistically, *Batf3*-deficient innate-like γδ T cells exhibit transcriptional changes that disrupt pathways governing actin cytoskeleton remodelling, immunological synapse organization and cellular identity. Notably, *Batf3*-deficient mice present decreased survival in a viral infection model highly dependent on innate-like γδ T cells. Together, our findings uncover a previously unrecognized BATF3-dependent pathway that controls γδ T cell morphology and function, profoundly impacting their biology.

## Introduction

T cells expressing the γδ T cell receptor (γδ TCR) represent a distinct lineage of lymphocytes with unique functional and developmental characteristics compared to their αβ T cell counterparts. Frequently described as unconventional lymphocytes, γδ T cells can adopt either innate- or adaptive-like characteristics, based on the type of thymic education they receive during development (1, 2). Unlike αβ T cells, which recognize peptide antigens presented by major histocompatibility complex (MHC) molecules, γδ T cells can respond to a broader range of antigens (3), including nonpeptidic small molecules, lipids, and stress-induced ligands. Moreover, innate-like γδ T cells can be activated directly by cytokines (4), and enabling them to act as first responders during inflammatory challenges across various tissues.

γδ T cells display substantial functional heterogeneity, which is closely linked to their specific subset identity; notably, the molecular composition of the γδ TCR strongly correlates with their functional fate (2). In mice, different γδ T cell subsets are defined by their TCR Vγ chain usage, displaying either adaptive- or innate-like features. Vγ5^+^ (nomenclature according to Heilig and Tonegawa (5)) dendritic epidermal T cells (DETCs) are exclusive to the skin epidermis and contribute to epithelial integrity and immune surveillance. These cells can also aid wound healing by producing IL-13, IFN-γ, and growth factors (6, 7). In the gut, Vγ7^+^ intraepithelial lymphocytes maintain barrier function and respond to pathogens by releasing TNF-α and Type I/III interferons (8). Vγ6^+^ cells are enriched in subepithelial sites like the skin, lung, uterus, gingiva, and meninges, producing IL-17A and IL-22, which are crucial for antimicrobial defense and tissue repair but can also promote autoinflammation (2, 9). By contrast, Vγ1^+^ and Vγ4^+^ subsets are more functionally diverse, found in lymphoid and non-lymphoid tissues, producing IL-17, IFN-γ, and IL-4. For example, Vγ4^+^ cells in the dermis are innate-like (10), but can also exhibit adaptive traits under inflammation (11). Similarly, Vγ1^+^ cells includes those that may clonally expand during infection (12) and others that phenocopy innate NKT-like cells (13).

In mice, innate-like γδ T cells exit the thymus in a functionally mature state, poised for cytokine production or cytotoxic activity (1, 9). Notably, this preprograming process is driven by TCR signal strength (14, 15) and transcriptional regulation, with key factors such as *Rorc* and *Tbx21* playing roles in guiding the differentiation of IL-17A- and IFN-γ-producing γδ T cells, respectively (16). Although several transcriptional regulators of γδ T cell identity, as judged by TCR usage and function, have been identified, our understanding of the full network controlling their development, function and homeostasis remains incomplete. In particular, there is no factor known to affect innate-like γδ T cells, irrespective of their function, e.g., IL-17 production or IFN-γ production.

Basic leucine transcriptional factor ATF-like (BATF) belongs to the family of activator protein-1 (AP-1) transcription factors, together with JUN, FOS, MAF among others (17). These transcription factors regulate numerous cellular processes, but BATF and the closely related molecule BATF3 rapidly emerged as unique regulators of the function and lineage commitment of different lymphoid and myeloid cell subsets (17). Notably, BATF3 has been identified as a key transcription factor in the development and function of conventional type 1 dendritic cells (cDC1s) (18) and has been widely studied in the context of dendritic cell biology (17). However, more recently, BATF3 has been shown to play critical roles in the generation of tissue-resident memory (T_RM_) CD8^+^ αβ T cells (19), tissue-resident CD4^+^ αβ T cells (20), and peripherally-generated regulatory T cells (21). However, its specific role in γδ T lymphocytes – the prototypical tissue-intrinsic T cells – has not yet been investigated.

Here we identify a precedent-setting role for BATF3 in regulating the biology of innate-like γδ T cell subsets irrespective of their functional phenoypes. Using *Batf3*^-/-^ mice, together with a mouse line specifically deficient in cDC1s (XCR1-DTA), we found that the absence of BATF3, but not cDC1s, led to a reduction of all innate-like γδ T cell subsets across multiple non-lymphoid tissues. Moreover, the remaining γδ T cells in *Batf3*^-/-^ mice presented pronounced functional and morphological defects. Mechanistically, BATF3 controls a gene network regulating cytoskeletal dynamics, cell activation, and subset differentiation acting after thymic maturation. Finally, BATF3 deficiency compromised resistance to influenza infection in neonatal mice, a model critically dependent on γδ T cell–mediated immunity. These findings reveal BATF3 as a key transcriptional regulator of γδ T cells and highlight this transcription factor as a potential target for modulating tissue-based immune responses in both health and disease.

## Results

### *Batf3*-deficient mice display a decrease in innate-like γδ T cells

To assess the impact of BATF3 deficiency on γδ T cell distribution across different tissues, we quantified γδ T cell number in lymphoid and non-lymphoid organs from *Batf3*^-/-^ mice and wild-type (WT) mice (**Fig. 1a**). The numbers of γδ T cells were decreased in *Batf3*^-/-^ animals in skin, gut and lung, but were unchanged in the epididymal white adipose tissue (eWAT), spleen and thymus (**Fig. 1a and 1b**). In the skin, we observed a clear reduction in DETC numbers (**Fig. 1c**); dermal γδ T cells presented a subtle but also significant decrease in numbers (**Fig. 1c**), indicating that BATF3 deficiency acts both on innate IFN-γ-skewed and IL-17-skewed γδ T cells. Furthermore, *Batf3*^-/-^ mice exhibited decreased numbers in both the Vγ4^+^ and Vγ6 (from here on defined as being Vγ1^-^Vγ4^-^Vγ5^-^) subsets of dermal γδ T cells (**Extended Data Fig. 1a**). We next assessed whether BATF3 influenced γδ T cell effector programming using CD44 and CD45RB – surface markers associated with thymic imprinting and functional polarisation (22). Lung γδ T cells from *Batf3*^-/-^ mice showed an increase in the proportion of CD44^+^CD45RB^+^ and CD44^-^CD45RB^+^ cells (subsets associated with IFN-γ production) at the expense of their CD44^hi^CD45RB^-^ counterparts (a phenotype associated with IL-17 production; **Fig. 1d**). Further analysis of lung-resident γδ T cell subsets, including Vγ1^+^, Vγ4^+^ and Vγ6 populations, revealed a significant decrease in their numbers in *Batf3*^-/-^ mice (**Extended Data Fig. 1b**). In the eWAT, while overall γδ T cell numbers were unaffected (**Fig. 1b**), there was a small but significant decrease in CD44^hi^CD45RB^-^ γδ T cells in *Batf3*^-/-^ compared to WT mice (**Fig. 1e**), along with a specific reduction in Vγ6 γδ T cells (**Extended data Fig. 1c**). We next examined secondary lymphoid organs. In the spleen, the total number of γδ T cells did not differ significantly between *Batf3*^-/-^ and WT mice (**Fig. 1a and 1b**). However, the proportion of CD44^hi^CD45RB^-^ γδ T cells was reduced in *Batf3*^-/-^ mice (**Fig. 1f**). These findings were mirrored by a specific decrease in Vγ6, but not Vγ1^+^ and Vγ4^+^, γδ T cells in the spleen (**Extended Data Fig. 1d**). However, we did not observe differences in the proportion of CD44^+^CD45RB^-^ γδ T cells (**Fig. 1g**) or the number of in Vγ6 γδ T cells in the thymus (**Extended Data Fig. 1e**). In summary, BATF3 deficiency results in a selective loss of innate-like γδ T cell subsets whose development depends on TCR signal strength (15), with the strongest effects observed in peripheral tissues such as the skin and lungs. These findings establish BATF3 as a key regulator of γδ T cell homeostasis and subset-specific distribution.

**Fig. 1.**
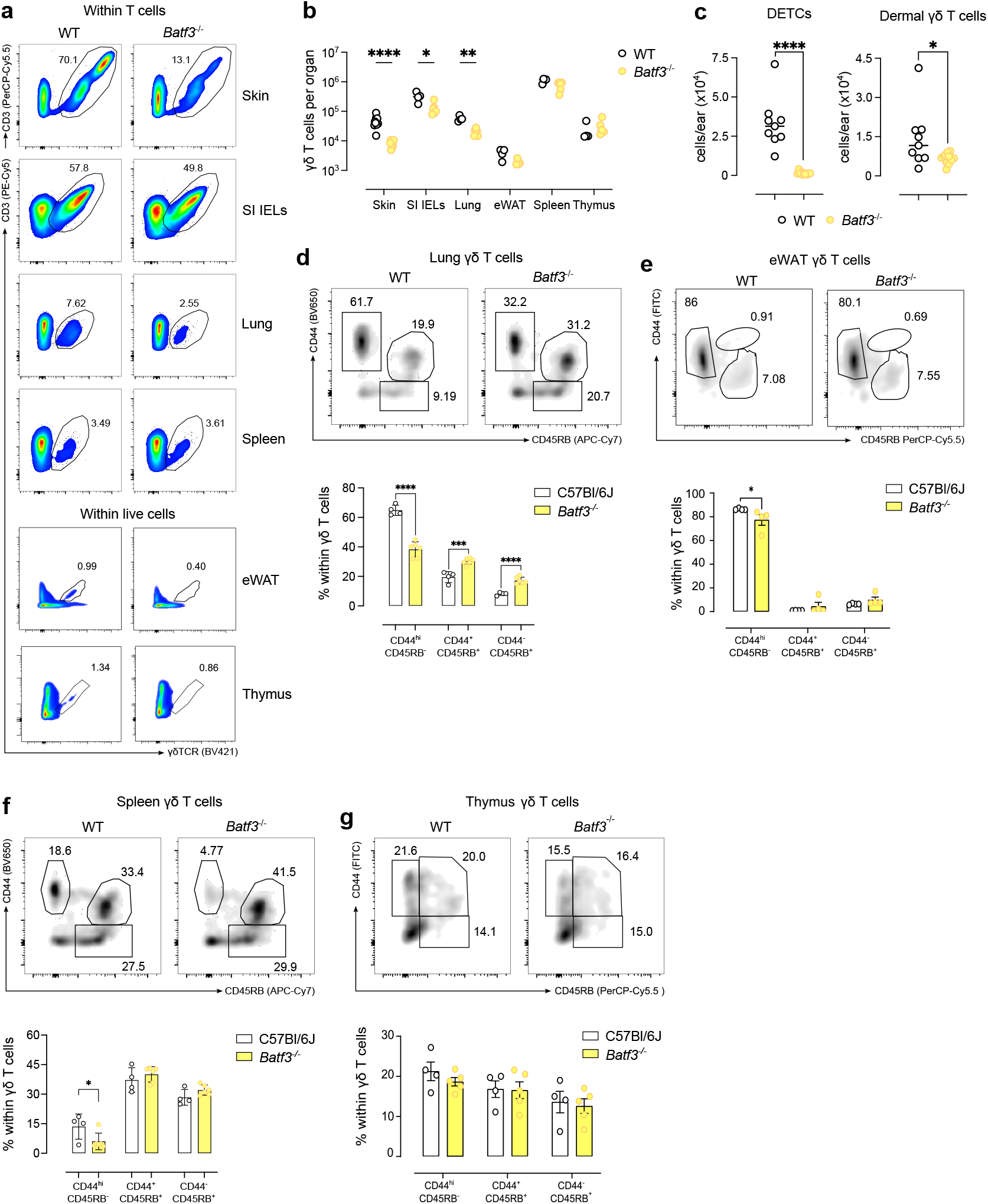
Altered distribution of γδ T cell subsets in *Batf3*-deficient mice. **(a)** Representative flow cytometry plots of γδ T cells within T cells in the ear skin, SI IELs, Lung and Spleen, and within CD45^+^ cells in eWAT, and thymus in WT and *Batf3*^-/-^ mice. **(b)** Quantification of γδ T cells per organ in WT and *Batf3*^-/-^ mice. **(c)** Quantification of DETCs and dermal γδ T cells (per ear) in WT and *Batf3*^-/-^ mice. **(d–g)** Representative plots (upper panels) and summary plots (lower/bottom panels) showing the frequencies of CD44^hi^CD45RB^-^, CD44^+^CD45RB^+^ and CD44^-^CD45RB^+^ subsets within γδ T cells in the lung **(d)**, eWAT **(e)**, spleen **(f)** and thymus **(g)** in WT and *Batf3*^-/-^ mice. **(b–c)** Data are presented as median (dots represent individual data points). **(d–g)** Data are presented as average, and error bars represent mean±SEM (dots represent individual data points). **(b–g)** Data are representative of at least 2 independent experiments. Normality of the samples was assessed, and statistical analysis was then performed accordingly. *, *p* < 0.05; **, *p* < 0.01; ***, *p* < 0.001; ****, *p* < 0.0001

### The effect of BATF3 on γδ T cells is not dependent on cDC1s

Since *Batf3*^-/-^ mice present a major defect in cDC1 development (18), we next asked whether BATF3 influenced γδ T cell homeostasis via cDC1. Hence, we analysed the distribution of total γδ T cells and their subsets across multiple tissues in XCR1-DTA^+^ mice – in which cDC1 cells are specifically depleted (23) (**Extended Data Fig. 2a**) but BATF3 expression in γδ T cells is preserved. Flow cytometry analysis revealed no significant decrease in the frequency or number of DETCs, dermal Vγ6 cells, or dermal Vγ4^+^ cells between XCR1-DTA^+^ and WT mice (**Fig. 2a**). In the lung, a similar result was observed, with no significant differences in the total number of γδ T cells or their subsets between XCR1-DTA^+^ and WT mice (**Fig. 2b**). Likewise, no significant differences were observed in the total number of γδ T cells in the thymus, nor in the numbers of Vγ4^+^ and Vγ6 thymic γδ T cells (**Fig. 2c**). Therefore, the absence of cDC1s does not explain the decreased γδ T cell numbers observed in *Batf3*^-/-^ mice, instead suggesting a cell-intrinsic role for BATF3 in regulating either γδ T cell development or peripheral maintenance.

**Fig. 2.**
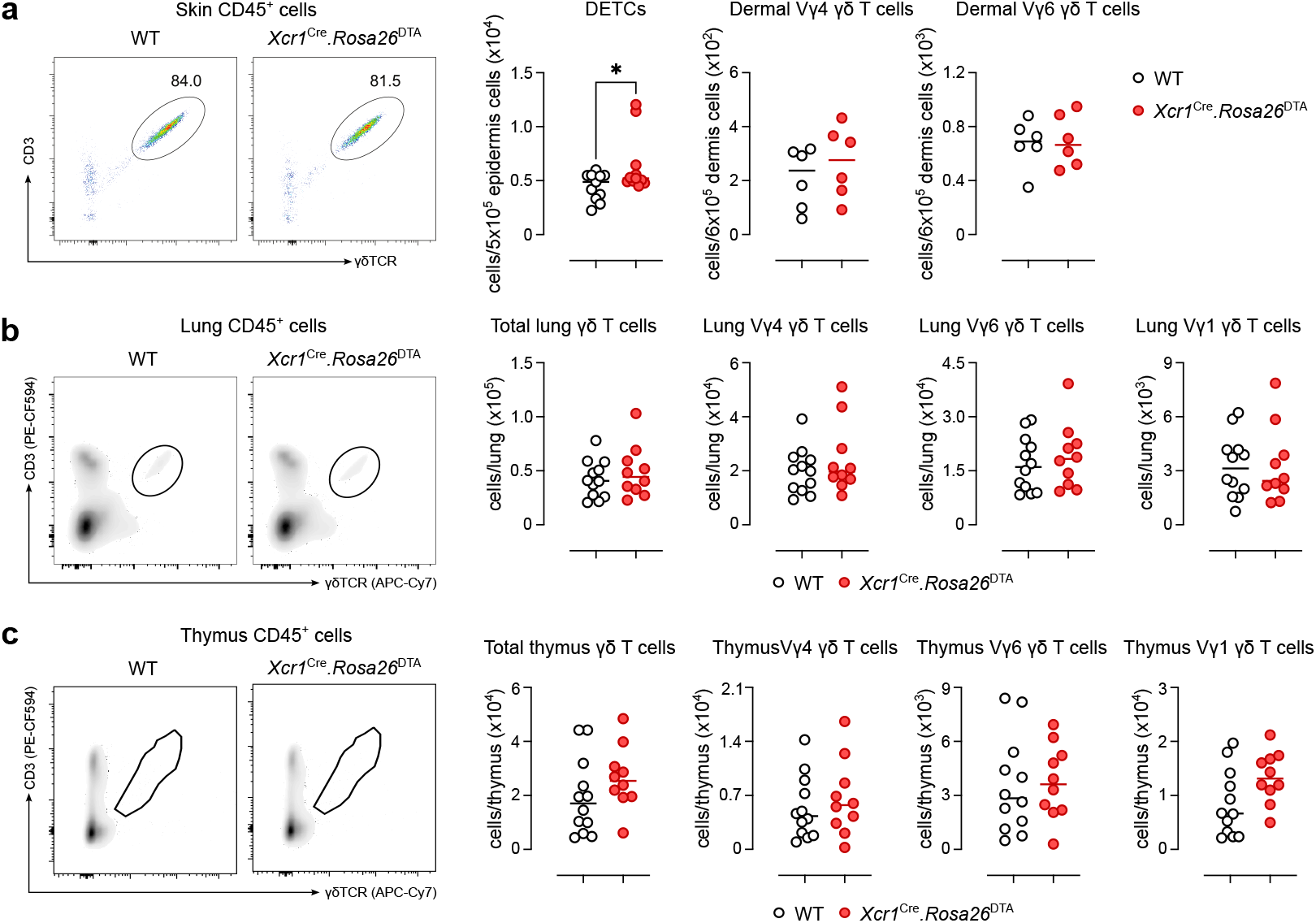
γδ T cell subset distribution in *Xcr1*^Cre^.*Rosa26*^DTA^ (XCR1-DTA) and WT mice. **(a–c)** Analysis of γδ T cell subsets in skin, lung and thymus in XCR1-DTA^+^ and WT mice. Representative flow cytometry plots of γδ T cells within CD45^+^ cells (left panels) and summary plots showing the number of total γδ T cells, DETCs, Vγ4, Vγ6 and Vγ1 γδ T cells (right panels) in the skin **(a)**, lung **(b)** and thymus **(c)**. Data are presented as median (dots represent individual data points). **(a–c)** Data representative of at least 2 independent experiments. Normality of the samples was assessed, and statistical analysis was then performed accordingly. *, *p* < 0.05

### BATF3 deficiency does not affect γδ T cell development in the embryonic thymus

To determine whether the observed defect in γδ T cells originates during their thymic development or arises in the periphery, we coupled the analysis of γδ T cell populations in the embryonic thymus with *in vitro* hanging drop foetal thymic organ cultures (HD-FTOC). Specifically, thymic cells from WT and *Batf3*^-/-^ mice were analysed at embryonic day (E) 16 to identify distinct γδ T cell populations based on the expression of Vγ5, Vγ1, and Vγ4, with the triple-negative population inferred to correspond primarily to Vγ6 cells. We observed no impaired development for the Vγ5^+^ and Vγ6 subsets in *Batf3*^-/-^ mice, when compared to their WT counterparts (**Fig. 3a**), although we did find a small but significant decrease in the frequency of embryonic Vγ1^+^ cells (**Fig. 3a**). Taken together, these results suggest that Batf3 deficiency does not impair thymic development of innate-like γδ T cells at this stage. Culturing of WT and *Batf3*^-/-^ FACS-sorted E15-16 thymocytes into E14-15 Rag1^-/-^ thymic lobes (**Fig. 3b**) showed comparable numbers of developing γδ T cells, including the Vγ5 and Vγ6 subsets, regardless of *Batf3* deficiency (Fig. 3c). Importantly, we observed no differences in the percentage of IL-17A^+^ and IFN-γ^+^ in mature γδ T cells in FTOC from WT and *Batf3*^-/-^ mice (**Extended Data Fig. 3a**). Thus, collectively our results suggest that BATF3 is dispensable for γδ T cell thymic development.

**Fig. 3.**
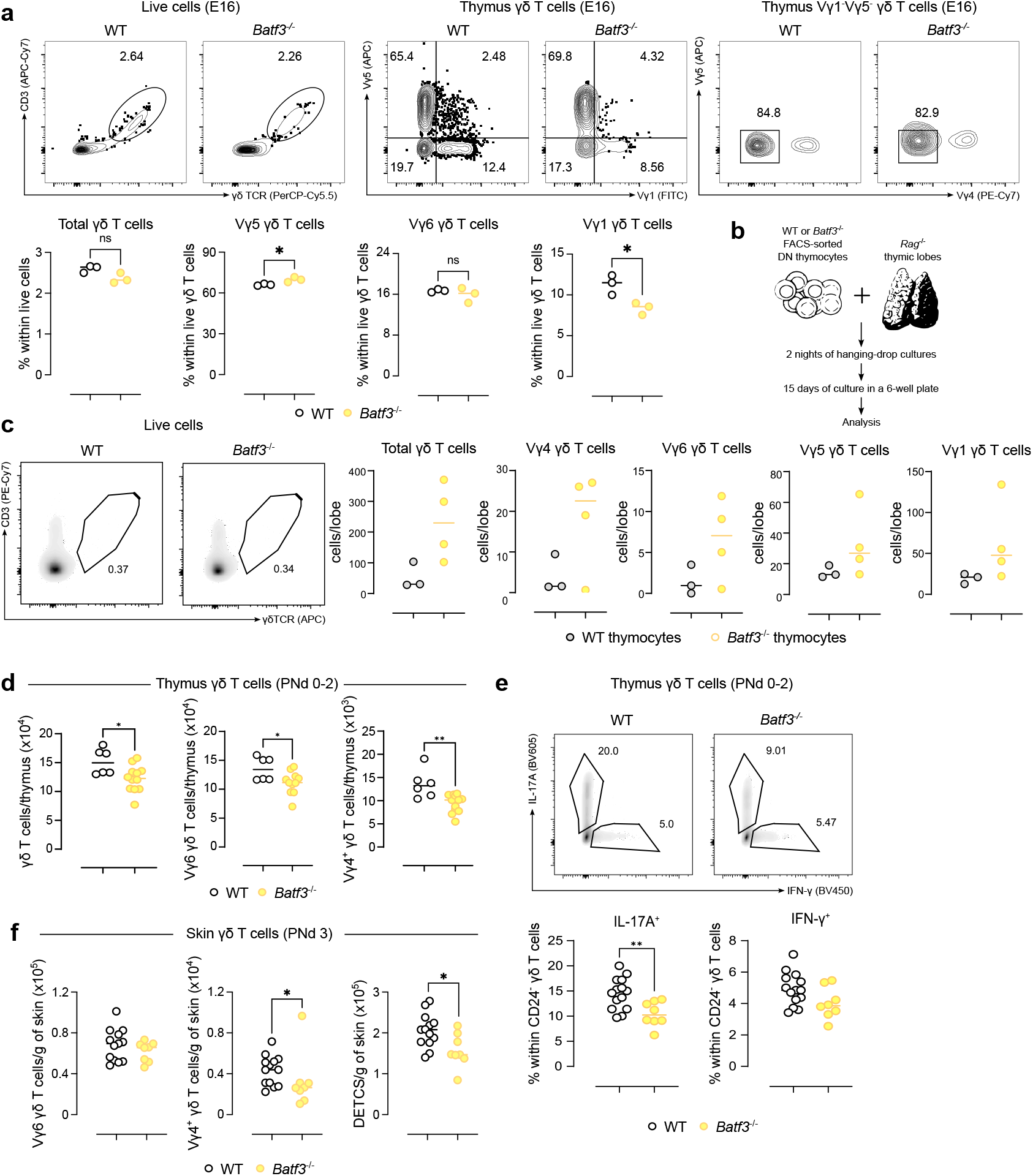
BATF3 deficiency does not affect γδ T cell development in the embryonic thymus. **(a)** Representative flow cytometry plots (left panels) and summary graphs (right panels) showing γδ T cells and Vγ subsets in embryonic day 16 (E16) thymus of WT and *Batf3*^-/-^ embryos. **(b)** Schematic showing the protocol utilised for generating the HD-FTOCs. **(c)** Representative flow cytometry plots and summary graphs showing total γδ T cells, Vγ5^+^, Vγ4^+^, Vγ1^+^ and Vγ6 subsets in HD-FTOCs of WT and *Batf3*^-/-^ mice. **(d)** Quantification of neonatal Vγ subsets in the thymus (PNd 0-2) of WT and *Batf3*^-/-^ mice. **(e)** Representative plots (upper panels) and individual data (bottom panels) showing the IL17-A and IFN-γ production by neonatal γδ T cells in the thymus of WT and *Batf3*^-/-^ mice after 4h of *ex-vivo* restimulation. **(f)** Quantification of neonatal γδ T cell subsets in the ear skin (PNd 3) of WT and *Batf3*^-/-^ mice. Data are presented as median (dots represent individual data points). Data in **(a, c)** is representative of 2 independent experiments. Data in **(d-f)** show a pool of 2 independent experiments. Normality of the samples was assessed, and statistical analysis was then performed accordingly. *, *p* < 0.05; **, *p* < 0.01; ****, *p* < 0.0001.

In order to pinpoint when BATF3 deficiency starts to impact γδ T cell homeostasis, we quantified total γδ T cells and individual subsets – including Vγ5^+^, Vγ1^+^, Vγ4^+^, and Vγ6 cells – in neonatal mice (post-natal day (PNd) 0-3). Evidently, neonatal thymic *Batf3*^-/-^ cells exhibited a significant reduction in the total number of γδ T cells, as well as in the Vγ4 and Vγ6 subpopulations, compared to WT mice (**Fig. 3d**). To assess the functional capacity of fully mature γδ T cells, we analysed cytokine production by CD24^-^ cells following *ex-vivo* stimulation. A marked decrease in IL-17A production was observed in *Batf3*^-/-^ mice compared to their WT counterparts (**Fig. 3e**), whereas IFN-γ production remained comparable between groups (**Fig. 3e, Extended Data Fig. 3b**), suggesting BATF3 starts to be crucial for innate-like γδ T cell survival or proliferation early postthymic development. Consistent with this observation, we also detected a reduction in the proportion of RORγt^+^PLZF^+^ cells within the CD44^hi^CD45RB^-^ γδ T cell subset in *Batf3*^-/-^ mice (**Extended Data Fig. 3c**). To rule out increased egress of γδ T cells from the neonatal thymus, we next examined the γδ T cell populations in the neonatal skin. *Batf3*^-/-^ mice exhibited a selective reduction in Vγ4^+^ dermal γδ T cells, while the Vγ6 population remained largely unaffected (**Fig. 3f**). In addition, *Batf3*^-/-^ animals displayed a significant decrease in both the total numbers of DETCs (Fig. 3f) and surface TCR expression (**Extended Data Fig. 3d**). Taken together, our findings indicate that *Batf3*^-/-^ γδ T cells develop normally in the thymus during foetal life, but their survival and maintenance is negatively impacted soon after birth, both in the thymus and in the periphery. Importantly, our data shows that BATF3 seems to be playing a role primarily in innate-like γδ T cell subsets.

### BATF3 regulates cytoskeleton dynamics and TCR signalling in DETCs

To explore the mechanisms by which BATF3 could be regulating γδ T cell homeostasis, we decided to focus on two major classes of innate-like γδ T cells impacted by *Batf3* deficiency: γδ intraepithelial lymphocytes (IELs; represented by DETCs and small intestine IELs) and innate IL-17 producers. For γδ IELs, we analysed the functional properties of the subsets most affected in *Batf3*-deficient mice. In the skin, flow cytometry revealed a marked reduction in the median fluorescence intensity (MFI) of CD3, γδTCR and Vγ5 in DETCs from *Batf3*^-/-^ mice compared to WT controls (**Fig. 4a**), further supporting a role for BATF3 in regulating γδ T cell function in the epidermis. Notably, this pattern extended to the other intraepithelial γδ T cell population, the gut-resident Vγ7^+^ γδ T cells, which also exhibited reduced MFI for both γδTCR and CD3 in the absence of BATF3 (**Extended Data Fig. 4a**). These findings suggest a broader role for BATF3 in supporting intraepithelial γδ T cell function across barrier tissues, through regulation of TCR complex expression and signalling capacity.

**Fig. 4.**
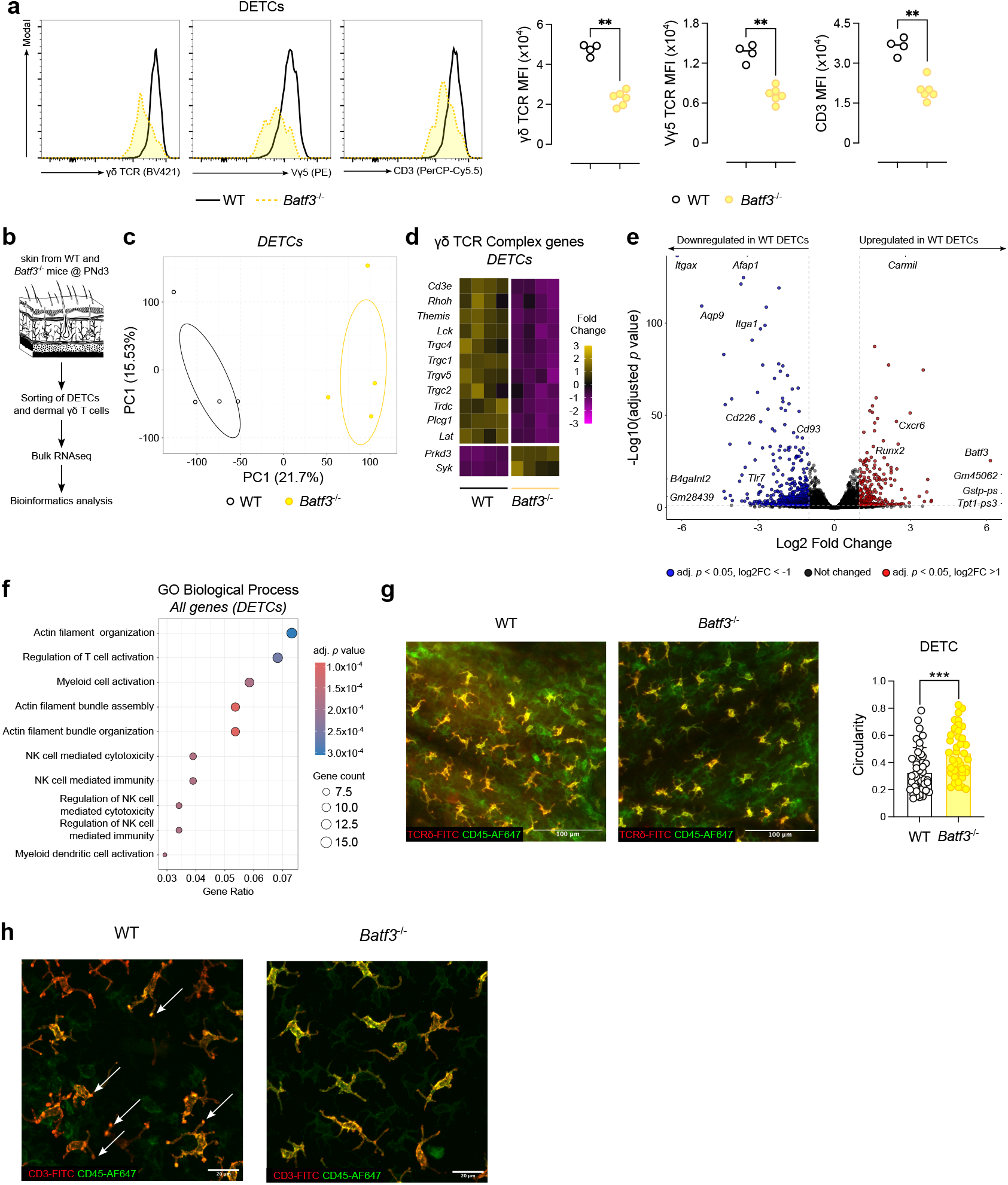
BATF3 regulates the morphology of DETCs. **(a)** Representative histograms (left panels) and summary plots (right panels) of TCR complex molecules median fluorescence intensity (MFI) in DETCs from WT and *Batf3*^-/-^ mice. **(b)** Experimental approach employed to obtain the transcriptional signature of DETCs and dermal γδ T cells from WT and *Batf3*^-/-^ mice (n = 4 mice per group) **(c)** Total sample variation in DETCs from WT and *Batf3*^-/-^ mice was summarised via PCA across all genes. **(d)** Normalised counts for genes in the γδ TCR Complex pathway converted to Z-scores. **(e)** DEGs from DETCs were visualised using a volcano plot, genes significantly upregulated (red), and downregulated (blue) were defined as having an adjusted *p* value of <0.05 and absolute log2 fold change of >1. **(f)** Overrepresented GO: biological processes in differentially regulated genes between DETCs from WT and *Batf3*^-/-^ mice. **(g)** Confocal microscopy of whole-mount ear skin preparations stained for TCRδ and CD45 in WT and *Batf3*^-/-^ mice (left and centre) [scale bars represent 100 µm]; quantification of DETC circularity in whole-mount skin preparations in WT (black) and *Batf3*^-/-^ (yellow) mice (right). **(h)** Confocal microscopy of whole-mount ear skin preparations stained for CD3 and CD45 in WT and *Batf3*^-/-^ [scale bars represent 20 µm]; white arrows indicate regions of concentrated CD3 staining (PALPs). **(a)** Data are presented as median (dots represent individual data points). Data in **(a, g-h)** are representative of at least 2 independent experiments. Normality of the samples was assessed, and statistical analysis was then performed accordingly. **, *p* < 0.01; ***, *p* < 0.001.

Next, we performed bulk RNA sequencing on FACS-sorted DETCs from neonatal WT and *Batf3*^-/-^ mice (and, thus overcoming potential issues with low cell numbers in the knockout mice; **Fig. 4b**). Striking differences in gene expression profiles were observed between DETCs from WT and *Batf3*^-/-^ mice, as illustrated by principal component analysis (PCA; **Fig. 4c**). As we had previously observed a consistent decrease in the expression of TCR complex molecules in *Batf3*^-/-^ DETCs (**Fig. 4a**), we investigated if genes controlling TCR signalling (**Extended Data Table 1**) were affected. Several TCR signalling-related genes, including *Lat, Rhoh, Themis, Lck, Cd3e* and multiple TCR structural genes were significantly downregulated in *Batf3*^-/-^ DETCs when compared to their WT counterparts, highlighting a disruption in pathways critical for T cell activation and signal transduction (**Fig. 4d**). On the other hand, genes such as *Prkd3* and *Syk* were upregulated in the absence of BATF3, potentially indicating compensatory mechanisms or shifts in TCR signalling dynamics (24) (**Fig. 4d**). An unbiased approach, however, painted a more complex picture. We identified more than 900 differentially expressed genes (DEGs) between WT and *Batf3*^-/-^ DETCs, significantly extending beyond TCR signalling pathways (**Fig. 4e and Extended Data Table 2**). Among the downregulated genes in *Batf3*^-/-^ DETCs were *Batf3* itself – serving both as an internal control, and reinforcing the idea that this transcription factor acts in a cell-intrinsic manner – as well as *Cxcr6* and *Runx2*, both being described as critical regulators of tissue residency (25, 26) (**Fig. 4e**). Conversely, in the absence of *Batf3*, several genes related to cell adhesion and innate response to pathogens (such as *Itgax, Itga1, Cd93, Cd226* and *Tlr7*) were found to be upregulated (**Fig. 4e**), suggesting a shift toward a more activated or migratory profile.

To further investigate the biological processes associated with the differentially expressed genes in *Batf3*^-/-^ mice, we performed Gene Ontology (GO) enrichment analysis of our dataset. As expected, pathways related to T cell activation and NK cell-mediated immunity were present in the top 10 enriched pathways among our DEGs (**Fig. 4f**). However, pathways involved in actin filament organization and actin filament bundle assembly were the most affected both by measures of statistical significance and the proportion of genes affected per pathway (Gene Ratio), indicating potential changes in cytoskeletal architecture and cell morphology in the absence of BATF3 (**Fig. 4f**). In summary, our transcriptional analysis permitted us to hypothesise that BATF3 orchestrates transcriptional networks governing TCR signalling, cell shape, and physical interactions with the microenvironment.

One defining characteristic of DETCs is their dendritic shape, recognised to be key in enabling tissue anchoring and TCR signalling focusing (27). Thus, to assess if the alterations in the transcriptional profile of *Batf3*^-/-^ DETCs also led to morphological alterations, we investigated DETC morphology by confocal microscopy in whole-mount epidermal preparations. In WT mice, DETCs appeared as highly dendritic, comprising the majority of CD45^bright^ cells in the tissue (**Fig. 4g**). In contrast, *Batf3*^-/-^ DETCs exhibited a markedly less dendritic morphology. Of note, CD45^dull^ cells, visible in an adjacent epidermal layer, correspond primarily to Langerhans cells and not T cells. To quantify these differences, we performed circularity analysis on epidermal γδ T cells. *Batf3*^-/-^ γδ T cells displayed significantly higher circularity values compared to WT, indicative of a more rounded morphology (**Fig. 4g**). When assessing the expression of the TCR complex-associated molecules CD45 and CD3, *Batf3*^-/-^ DETCs exhibited a distinct surface expression profile compared to WT, with reduced co-localization of CD3 and phosphorylated tyrosine-rich aggregates on cellular projections (PALPs) (27), which were nearly absent in *Batf3*^-/-^ cells (Fig. 4h).

Altogether, our data indicate that *Batf3* deficiency in DETCs results in substantial changes in gene expression, cytoskeletal organization, cell morphology, and surface receptor distribution, underscoring its critical role in the maintenance and functional integrity of epidermal γδ T cells (and small intestine IELs).

### BATF3 regulates γδ17 T cell identity and function

To explore the impact of *Batf3* deficiency on the transcriptional regulation of γδ17 T cells, we performed bulk RNA sequencing on FACS-sorted dermal γδ T cells from neonatal WT and *Batf3*^-/-^ mice (**Fig. 4b and Fig. 5a**). Akin to what we observed with DETCs (**Fig. 4c**), *Batf3*^-/-^ and WT dermal γδ T cells showed a markedly distinct transcriptional profile (**Fig. 5a**), with more than 400 DEGs between *Batf3*-deficient and -sufficient cells (**Fig. 5b and Extended Data Table 3**). Importantly, *Batf3* was among the top upregulated genes in WT dermal γδ T cells when compared to their knockout counterparts (**Fig. 5b**), confirming that this transcription factor is also expressed in γδ17 T cells. Interestingly, when we looked at the core genes composing the γδ17 T cell transcriptional signature, we found most of the genes, including *Rorc, Sox13* and *Blk*, to be significantly downregulated in *Batf3*^-/-^ dermal γδ T cells (**Fig. 5c**). Notable exceptions were the genes coding for cytokines, such as *Il17a, Il22, Ifng* and *Csf1*, which were all upregulated in *Batf3*^-/-^ dermal γδ T cells (**Fig. 5c and Extended Data Table 4**). Furthermore, the collection of genes that are significantly upregulated in WT dermal γδ T cells positively correlated with T cell identities extracted from the PanglaoDB database for scRNAseq studies (**Extended Data Fig. 5a**); genes upregulated in *Batf3*^-/-^ dermal γδ T cells, on the other hand, did not match T cell identities (**Extended Data Fig. 5b**). Overall, transcriptomic profiling of dermal γδ T cells from *Batf3*^-/-^ mice revealed marked alterations in gene expression, ultimately impacting on the establishment of their γδ17 T cell transcriptional signature, suggesting an essential role for BATF3 in coordinating immune cell function.

**Fig. 5.**
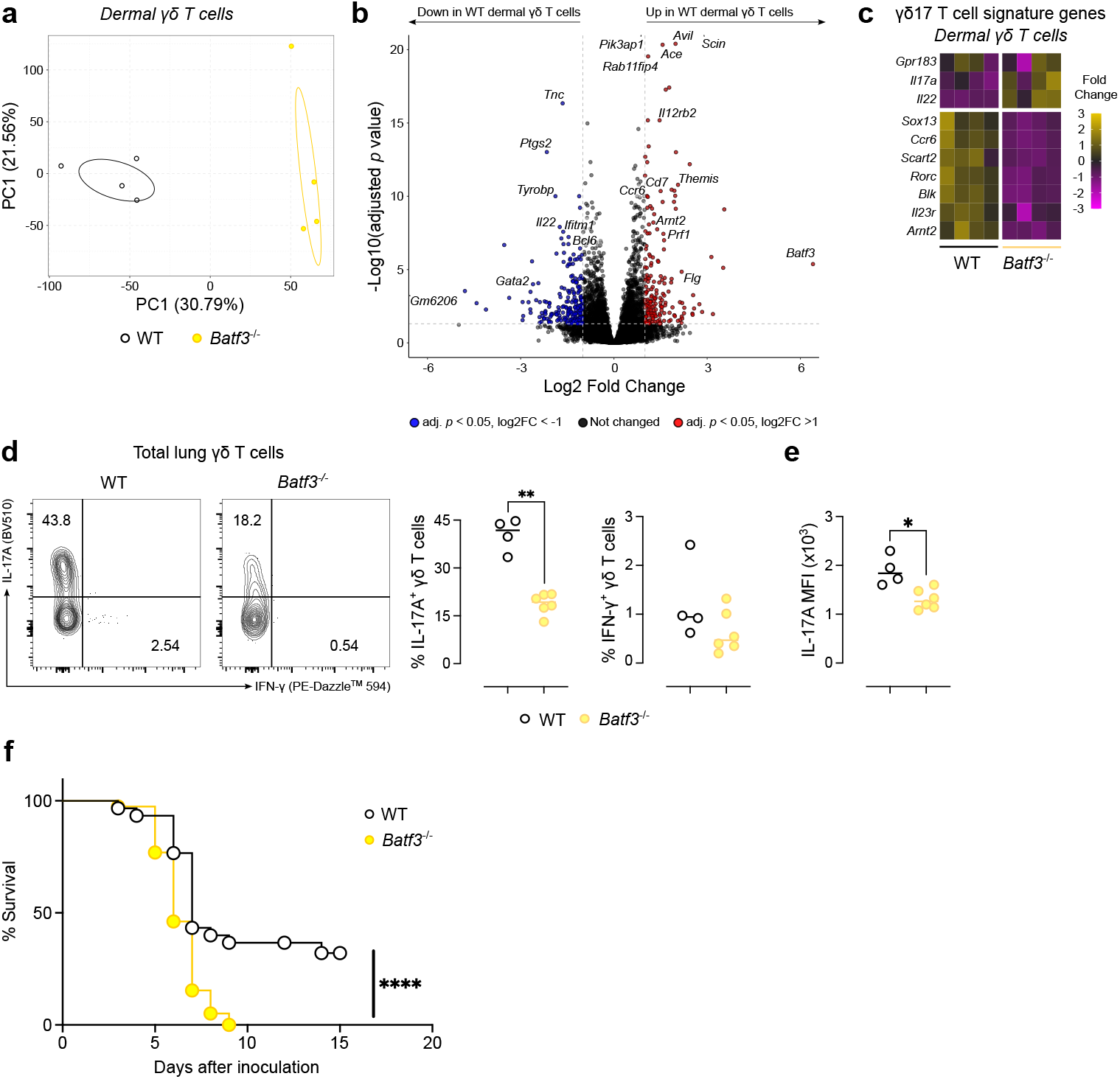
Increased susceptibility of *Batf3*^-/-^ neonatal mice to influenza virus infection. **(a)** Total sample variation in dermal γδ T cells from WT and *Batf3*^-/-^ mice was summarised via PCA across all genes (n = 4 mice per group). **(b)** DEGs from dermal γδ T cells were visualised using a volcano plot, genes significantly upregulated (red), and downregulated (blue) were defined as having an adjusted *p* value of <0.05 and absolute log2 fold change of >1. **(c)** Normalised counts for γδ17 T cell signature genes from WT and *Batf3*^-/-^ mice converted to Z-scores. **(d)** Representative flow cytometry plots (left panels) and summary plots (right panels) depicting IL-17A and IFN-γ production by γδ T cells in the lung after 4h of *ex-vivo* stimulation comparing WT and *Batf3*^-/-^ mice. Data are presented as median (dots represent individual data points). **(e)** Median fluorescence intensity (MFI) expression of IL-17A in γδ T cells in the lung of WT and *Batf3*-deficient mice shown in **(d). (f)** Survival of 6-to 7-day-old WT and *Batf3*^-/-^ neonatal mice following intranasal infection A/HKx31 (H3N2) influenza virus. **(d-e)** Data representative of at least 2 independent experiments. **(f)** Data show a pool of four experiments. Normality of the samples was assessed, and statistical analysis was then performed accordingly. *, *p* < 0.05; **, *p* < 0.01; ****, *p* < 0.0001.

Next, we assessed the cytokine-producing capacity of *Batf3*-deficient γδ T cells. In the lung, in a stark contrast to the transcriptomic data (**Fig. 5c**), IL-17A production by γδ T cells was significantly impaired, while IFN-γ production remained unaffected (**Fig. 5d**). This was reflected by a marked reduction in both the percentage of IL-17A^+^ γδ T cells and their cytokine expression levels (**Fig. 5d and 5e**). In order to determine whether the reduction in IL-17A-producing γδ T cells was tissue-specific or part of a broader phenotype, we further analysed γδ T cells in eWAT. Consistent with our findings in the lung, IL-17A production was significantly reduced in *Batf3*^-/-^ mice, supporting a more generalized impairment in this effector function (**Extended Data Fig. 5c and 5d**); CD27^+^ type 1 γδ T cells, on the other hand, were found to be increased in *Batf3*-deficient mice (**Extended Data Fig. 5c**). This apparent disconnect between the IL-17A mRNA and protein levels in *Batf3*^-/-^ mice may reflect a dysregulated activation state: in the absence of BATF3, γδ T cells may receive incomplete TCR or co-stimulatory signals, leading to impaired expansion and maintenance but a skewed transcriptional profile. Such a phenotype is consistent with a stress or dysregulated signalling response, where cells fail to sustain proper effector functions yet transiently increase cytokine gene expression.

To investigate the physiological relevance of *Batf3* deletion and consequent impairment of γδ17 T cell function, we assessed the impact of *Batf3* deficiency during neonatal influenza infection, a model where survival is tightly linked to the production of IL-17 by γδ T cells (28). Neonatal WT and *Batf3*^-/-^ mice (aged 6 to 7 days) were infected with A/HKx31 (H3N2) influenza virus, and survival was monitored over time (**Fig. 5f**). *Batf3*^-/-^ mice exhibited a significant reduction in survival compared to WT controls, with the most pronounced differences observed at later time points post-infection (days 8–10) (**Fig. 5f**). These findings suggest that the reduced number of γδ17 T cells in *Batf3*-deficient neonates (and their impaired capacity to produce IL-17A) may compromise host defence and contribute to increased susceptibility to influenza infection. This result highlights the pathophysiological alterations due to the impact of *Batf3* deficiency in γδ17 T cells.

## Discussion

The γδ T cell compartment plays a central role in maintaining tissue integrity and in orchestrating rapid responses to infection and stress. These cells are enriched at epithelial surfaces and mucosal barriers, where their functions range from cytokine production to tissue repair and immune surveillance (2). While recent studies have begun to uncover the heterogeneity and plasticity of γδ T cells, the molecular mechanisms that sustain their tissue residency and long-term function remain incompletely understood. In this study, we identify BATF3 as a critical, non-redundant, regulator of innate-like γδ T cells, including epidermal DETCs and Vγ4^+^ and Vγ6 subsets. BATF3, originally characterized for its indispensable role in the development and function of cDC1s (17, 18) γδ T cell landscape. Mice lacking *Batf3* exhibit a profound reduction in tissue-resident γδ T cells, accompanied by transcriptional, phenotypic, and morphological alterations.

DETCs have been shown to closely interact with cDC1s, via a XCL1-XCR1 axis that regulates their positioning during viral infection (29). Moreover, cDC1s have been shown to control IL-17 production by CD4^+^ T cells both during viral and fungal infection (30, 31). Thus, it was conceivable that the marked decrease in innate-like γδ T cells observed in *Batf3*^-/-^ mice stemmed from the disruption of key cell-cell interactions between cDC1s and γδ T cells. However, by employing XCR1-DTA^+^ mice, and selectively ablating cDC1s, we showed that the numbers and distribution of γδ T cells, including epidermal DETCs and dermal Vγ4^+^ and Vγ6 subsets, as well as thymic γδ T cells, remained unaltered. Thus, our findings demonstrate that the impaired survival of innate-like γδ T cell subsets observed in *Batf3*^-/-^ mice is not a secondary consequence of cDC1 depletion but likely reflects a γδ T cell-intrinsic defect. Importantly, the detection of *Batf3* expression in γδ T cells adds strength to this hypothesis.

Our results align with emerging literature highlighting BATF3 as a broader regulator of tissue-resident and memory T cell programs. In CD8^+^ T cells, BATF3 is required for the maintenance of memory populations (19). In addition, *Batf3* deficiency impairs the generation of a CXCR6^+^ population of lung-resident CD4^+^ T cells. More recently, CRISPR screens in human T cells have identified BATF3 as a top regulator of memory differentiation and exhaustion programs, with BATF3 overexpression improving CAR-T cell function and persistence (32). Our findings extend this functional profile to innate-like γδ T cells, which are often tissue-resident and display features of memory, such as high CD44 expression and rapid effector cytokine production (33).

In cDC1s, BATF3 interacts with *Irf8* during development to reinforce lineage commitment (34–36). In CD8^+^ T cells, transient BATF3 expression following priming, suppresses the proapoptotic molecule BIM, partially explaining how BATF3 regulates immune memory formation (19). Our data indicate that BATF3 mainly impacts the survival of γδ T cells in the periphery rather than mediating their thymic development. An exception is the observed decrease in γδ T cells in the neonatal thymus. However, as Batf3^-/-^ γδ T cells develop normally during embryonic life and in hdFTOCs, a decreased γδ T cell peripheral survival could alter neonatal thymic output through a feedback mechanism. Unexpectedly, we found transcriptional and morphological defects in Batf3-deficient DETCs, suggesting that BATF3 regulates cytoskeleton dynamics. Remarkably, DETCs are highly dependent on their dendritic morphology for continuous environmental surveillance, including recognition of stress signals and TCR-mediated activation. This unique morphology is underpinned by complex cytoskeletal organization and actin remodelling, which are essential for forming peripheral TCR-enriched microdomains (PALPs) (27, 37). In Batf3^-/-^ mice,DETCs displayed significant morphological alterations, suggesting that BATF3 may control transcriptional programs related to cytoskeletal architecture. Notably, CD8^+^ T_RM_ cells also rely on cytoskeletal remodelling for tissue retention and function (38), suggesting that BATF3 may regulate a shared module of tissue residency across T cell lineages.

Although our results do not explain how BATF3 sustains γδ17 T cell lineage commitment, impairment of immune synapse formation and disruption of TCR signalling could also explain the marked decrease of γδ17 T cells in *Batf3*^-/-^ mice. Despite increased *Il17a* mRNA expression in dermal γδ T cells from *Batf3*-deficient mice, their cytokine production is reduced upon restimulation. This suggests a disconnection between transcription and functional output, possibly due to disrupted translation, post-transcriptional control, or defective signal transduction. These findings underscore BATF3’s role in coordinating not only identity-defining transcriptional programs but also the downstream machinery required for optimal effector function. To explore the functional implications of BATF3 deficiency, we assessed susceptibility to influenza infection during the neonatal period—a model in which survival depends critically on IL-17 production by lung-resident γδ T cells (28). *Batf3*^-/-^ neonates displayed significantly reduced survival following influenza infection, particularly at late time points (days 8–10), correlating with the impaired γδ17 T cell pool. These results strongly support a functional requirement for BATF3 in maintaining protective γδ17 T cell-mediated immunity during neonatal infection.

Altogether, our study uncovers a novel role for BATF3 in controlling tissue-resident γδ T cell biology, independent of its role in dendritic cell development. These findings expand the functional repertoire of BATF3 and underscore its broader relevance across immune lineages. Importantly, our data – alongside recent studies – highlight the need for caution when using *Batf3*^-/-^ mice to study cDC1s, as BATF3 also regulates T cell function. Future studies will benefit from conditional approaches that allow cell-type–specific manipulation of BATF3 to fully dissect its multifaceted roles in immune regulation.

## ACKNOWLEDGEMENTS

We thank the staff at the Biological Services Facility and the Flow Cytometry Facility – in particular David Chapman – at The University of Manchester and the CNIC and UCM facilities and personnel for their technical assistance. We thank Matt Hepworth, Joanne Konkel, Grace Mallet and Rita Domingues for helpful discussions, sharing of resources and technical advice. This work was funded by the Wellcome Trust (106898/A/15/Z) and MRC (MR/V011235/1) to J.E.A., European Molecular Biology Organization (LTF 191-2019) and European Commission Marie Sklodowska-Curie Individual Fellowship (ref. 101025781) to P.H.P.; M.M-R was supported by funding from the Agencia Española de Investigación (AEI) (PID2022-139095OA-I00 and RYC2021-034348-I) and Fundación Fero (IV Ayuda de Investigación en Cáncer FERO-ASEICA). A.Z-C. was supported by an AEI predoctoral fellowship (PREP2022-000845). R.G.R.P. and B.S.-S. were supported by “la Caixa” Foundation under project LCF/PR/HR24/00929. S.I. was supported by a PID2021-125415OB-I00 grant from the Agencia Estatal de Investigación (AEI), co-funded by FEDER, UE. Fellowships: E.H.G.: FPI fellowship (PRE2019-087509) from the Spanish Ministry of Science, Innovation, and Universities. A.R.U. is supported by “Ayuda para la contratación de personal investigador predoctoral en formación 2022 (PIPF-2022/SAL-GL-24581) from the Community of Madrid.

## AUTHOR CONTRIBUTIONS

Conceptualization: P.H.P., E.H.-G., S.I. and M.M.-R.; Methodology and Investigation: P.H.P., E.H.-G., J.E.P., M.M.-R., A.Z-C, A.R-U.: R.G.R.P.; Data Analysis: P.H.P., E.H.-G., A.R-U.; J.E.P., M.M.-R.; Resources: A.S.M., C.R. e S., J.E.A., S.I., D.S.,B.S.-S.; Writing – original draft: P.H.P., E.H.-G., S.I. and M.M.-R.; Writing – review and editing: P.H.P., E.H.-G., B.S.-S., S.I. and M.M.-R.

## Methods

### A. Mice

*Batf3*-deficient, XCR1-DTA^+^ and WT control mice were bred and maintained under specific pathogen-free conditions. Mice were housed in groups of 2–5 animals per cage and maintained on a C57BL/6 background. A mix of *Batf3*^+/+^ littermates, XCR1-DTA^-^ mice, and C57Bl6/J were used as WT controls. In the case of XCR1-DTA^+^ mice, control mice included *Xcr1*^Cre^ mice expressing CRE but not DTA (CRE^+^ DTA^-^), and vice versa, mice expressing DTA but not CRE (CRE^-^ DTA^+^). All experiments were performed in compliance with institutional guidelines, and most experiments were repeated across the three main sites involved in this study (The University of Manchester, Francis Crick Institute and CNIC). All animal studies were approved by the local ethics committee, adhering to EU Directive 86/609/EEC and Recommendation 2007/526/EC on the protection of animals used for scientific purposes. These regulations are enforced in Spanish law under Real Decreto 1201/2005. In the UK, Experiments were performed in accordance with the United Kingdom Animals (Scientific Procedures) Act of 1986 (project license number 70/8547 and PP4115856).

### B. Organ harvesting

Mice were euthanized either by intraperitoneal (i.p.) injection of 200 mg/kg sodium pentobarbital (DOLETHAL, Vetoquinol) or by asphyxiation in a rising concentration of CO2. Prior to harvest of the organ of interest, in some experiments the mouse heart was perfused with 10 ml of PBS.

#### eWAT

eWAT samples were stored at 4 °C in 2 ml of HBSS without calcium or magnesium (Thermo Fisher Scientific) supplemented with 0.5% BSA (Sigma). Cell isolation and flow cytometry analysis: The stromal vascular fraction (SVF) cells in eWAT were isolated as previously described (39). In brief, after being mechanically disaggregated, eWAT was digested in HBSS supplemented with 0.5% BSA containing 4 mg/ml type II collagenase (Worthington Biochemical Corporation) for 20 min at 37 °C. The digestion was stopped with FBS. The cells were filtered twice through 100 µm (Fisher Scientific) and 30 µm (BD Biosciences) cell strainers and centrifuged sequentially at 600 g for 10 min and 5 min. The pellet was diluted in FACS buffer (PBS supplemented with 2,5% FBS, 2 mM EDTA, and 0.01% sodium azide) before proceeding with antibody staining.

#### Thymus

Thymi were harvested in PBS and then filtered through 100 µm cell strainer (Fisher Scientific) using a syringe plunger. Then the strainer was rinsed with ice-cold FACS buffer to wash the remaining cells and used for further analysis.

#### Lung

Lungs were harvested in HBSS and digested with 250 µg/ml Liberase TL (Sigma Aldrich) and 50 µg/ml DNase I (Sigma Aldrich) in HBSS for 30 minutes at 37 ºC. After the incubation, the tissue was passed 3 times through an 18-gauge syringe and incubated again another 15 minutes at 37ºC. Digestion was stopped by adding 10% of heat inactivated Fetal Bovine Serum (FBS) and placing the samples on ice. Lungs were passed 3 times through an 18-gauge syringe and filtered through 100 µm cell strainer (Fisher Scientific) using a syringe plunger. Then the strainer was rinsed with icecold FACS buffer to wash the remaining cells. Tissue homogenates were filtered through a 30 µm cell strainer (BD Biosciences) and used for further analysis.

#### Spleen

Spleen were harvested in HBSS and mechanically dissociated using a syringe plunger. Tissue homogenates were filtered through a 70 µm cell strainer (Fisher Scientific). Then cell pellets were lysed using Red Blood Cell Lysing buffer (Sigma) for 5 minutes at room temperature. Then the cells were washed with FACS buffer and used for further analysis. *Skin* Ears were dissected, and the two ear sheets were split using a pair of forceps. Split ear sheets were then digested with Liberase (0.25 mg/mL; Roche) and DNase I (0.5 mg/mL; Sigma-Aldrich) in 0.5mL RPMI 1640 at 37°C, 200 rpm, for 1h 45 mins. The supernatant and the remainder of the tissue were added to a Medicon (BD Biosciences) then homogenised with the help of a Medimachine for 5 minutes. The resulting tissue homogenate was then passed through a 70-µm cell strainer and washed in 10mL of RPMI 1640 containing 10% Fetal Bovine Serum (FBS).

### C. Flow Cytometry

Single-cell suspensions were prepared from the organs specified above. Cells were stained with antibodies against γδTCR, CD3, and subset-specific markers for flow cytometric analysis. Samples for Flow cytometry were incubated with LIVE/DEAD™ Fixable Blue Dead Cell Staining Kit, for UV Light Excitation or for 405 nm excitation (ThermoFisher) in PBS to exclude dead cells. After washing with PBS, samples for flow cytometry analysis were stained in FACS Buffer. Cells were preincubated for 10 min at 4 °C with anti-mouse CD16/CD32 (clone 2.4G2, Tonbo Bioscience, or BD Biosciences) before staining with the appropriate antibody cocktails. Events were acquired on FACSymphony or FACSFortessa (both BD Biosciences), and data were analyzed using FlowJo software (TreeStar). For intracellular staining, cells were fixed and permeabilized with a Foxp3/ Transcription Factor Staining Buffer Kit (Tonbo Bioscience, or Invitrogen) or fixed in 4% PFA and intracellularly stained during permeabilization with 0.1% saponin for intracellular cytokine analysis. Each experiment contained a minimum of three biological replicates, and a minimum of two independent experiments was performed. Intracellular cytokine staining was performed after stimulation with PMA (50 ng/mL) and ionomycin (1 µg/mL) during 4 hours, with brefeldin A (5 µg/mL) and BD GolgiStop™ (following the manufacturer’s recommendation) were added to the culture for the last 2 hours. Alternatively, cells were incubated in a 96 round-bottom wells plate in a 2-3×10^6^ cells/well concentration with a cell stimulation cocktail containing protein transport inhibitor (eBioscience) for 3h to 4h at 37 °C. In both cases intracellular staining was carried out as described above.

### D. Cell sorting

For cell sorting of dermal γδ T cells and DETCs, skin tissue was digested as described above and, additionally, single cells were isolated by passing the tissue through a 40-µm cell strainer, followed by a 70% / 30% Percoll (Sigma-Aldrich) gradient and 30-min centrifugation at 2400 rpm. Leukocytes were recovered from the interface, resuspended, and immunostained as described above. Dermal γδ T cells were defined as CD45^+^CD64^-^CD11b^-^CD11c^-^MHC-II^-^FcεR1^-^Thy1.2^+^CD3^+^γδTCR^+^Vγ5^-^, DETCs were defined as CD45^+^CD64^-^CD11b^-^CD11c^-^MHC-II^-^FcεR1^-^Thy1.2^+^CD3^+^γδTCR^+^Vγ5^+^ and then electronically sorted on a FACSAria (BD Biosciences) into sterile RNase-free Eppendorfs containing 200µL of lysis buffer (Qiagen), for posterior mRNA isolation; each mouse was treated as one individual sample.

### E. FTOCs

#### Timed Matings

RagKO male and female mice served to generate host lobes, while wild-type (WT) and *Batf3*-deficient mice served as sources for donor cells. Timed matings for both host and donor embryos were scheduled in a staggered manner as needed to ensure appropriate developmental stages. For host lobes, embryos were collected at embryonic day (E) 14–15, whereas donor lobes were obtained at E15–16. Firstly, male breeders were housed individually in their cages prior to matings. For increased success, we placed the male bedding in the female cages and the female bedding in the male cages 2 days prior to scheduled matings. The mice were left to breed 2 nights. Then, the males were removed from the female cages.

#### Hanging drop FTOC Fetal thymic organ culture (HD-FTOCs)

We harvested the thymus from the embryos at embryonic days 15-16 and double negative (DN) thymocytes were sorted from the thymus of WT and *Batf3*-deficient mice. Then, we added 20,000-10,000 DN cells to each RagKO lobe in a Terasaki plate and we incubated the hanging-drop cultures at 37 ºC for 2 nights until the lobes were seeded with cells from the cell suspension. After 2 nights, we removed the Terasaski plate from the incubator and inverted it. We collected the repopulated lobes and we transfered them to the filters in a plate with media. We removed the media every 3-4 days, and we cultured HD-FTOCs for 11-13 days to allow thymocyte development. Finally, thymus was processed as described above.

### F. Bulk RNA sequencing

RNA was isolated from sorted skin cells using the Single Cell RNA Purification Kit (Norgen). Samples were sent to Francis Crick Insitute where QC checks and sequencing were performed. In brief, cDNA was prepared from 10 ng input RNA using the NuGEN Ovation RNA-Seq System (V2; Manchester, United Kingdom), and libraries constructed using the NuGEN Ultralow Library System V2. Both steps followed the manufacturer’s instructions. The resulting libraries were pooled for sequencing on an Illumina HiSeq 2500 platform (San Diego, Calif) with singleended 75-bp reads.FASTQ files were received, which were then aligned to GENCODE mouse genome (version M33, Jan 2020), and quantified using Salmon software (v1.10.3) using –validatemappings and –gcBias flags (COMBINE-lab). Annotated and quantified data was then imported into RStudio (v2022.07.2+576) using the Tximeta R package (v4.2.3). Genes with an average raw count of less than ten across all groups were removed. Principal component analysis (PCA) was used to summarise total variation within the dataset. Differential expression analysis was performed using the DE-Seq2 R package (v1.42.0) and volcano plots and dot plots were then generated using ggplot2 R package (v3.5.1). The networks and functional analyses of DEGs were generated either via the clusterProfiler (v4.10.1) and enrichplot (v1.22.0) packages or via import into PANTHER (v19.0) via the web GUI (https://geneontology.org).

### G. Immunofluorescence Staining

Whole depilated splitear sheets were flattened in histology cassettes, fixed in 10% neutral-buffered formalin for 2 h at room temperature (RT), and permeabilized in 0.5% Triton X-100 for at least 1 h at RT. Epidermal sheets were blocked in PBS containing 5% normal goat serum (NGS) and then stained for 1 h at RT with antibodies diluted in PBS containing 5% NGS and 0.5% BSA. Z-stacks were acquired on a Leica SP5 confocal microscope (Leica Microsystems, Deerfield, IL) using a 10×/1.25 NA objective, and images were processed and analysed in Fiji. The morphological parameter sphericity was calculated as the ratio of cell-object border length to its volume, ranging from 0 to 1, with higher values corresponding to more spherical objects.

### H. Viral infections

Neonatal WT and *Batf3*^-/-^ mice (aged 6 to 7 days) were infected with A/HKx31 (H3N2) influenza virus. Neonatal mice were infected with a dose of 50 or 75 PFU per 3 grams of body weight, depending on their age (6 or 7 days old, respectively) and survival was monitored over time.

### I. Statistical Analysis

1. Statistical analysis was performed using GraphPad Prism 8 (GraphPad Software). Statistical significance was determined using Student’s t-test or ANOVA, as appropriate. Statistical significance for comparison between two sample groups coming from a normal distribution was determined by parametric two-tailed unpaired Student’s t-test or Mann & Whitney’s test for non-normal distributed samples. Normality was determined by comparison of the Anderson-Darling, D’Agostino-Pearson omnibus, Shapiro-Wilk and Kolmogorov-Smirnov with Dallal-Wilkinson-Lillie approximation normality tests. Survival between the different groups was analyzed by Long-rank (Mantel-Cox) test. All differences with *p* values < 0.05 were considered significant.

## Extended Data

**Extended Data Figure 1.**
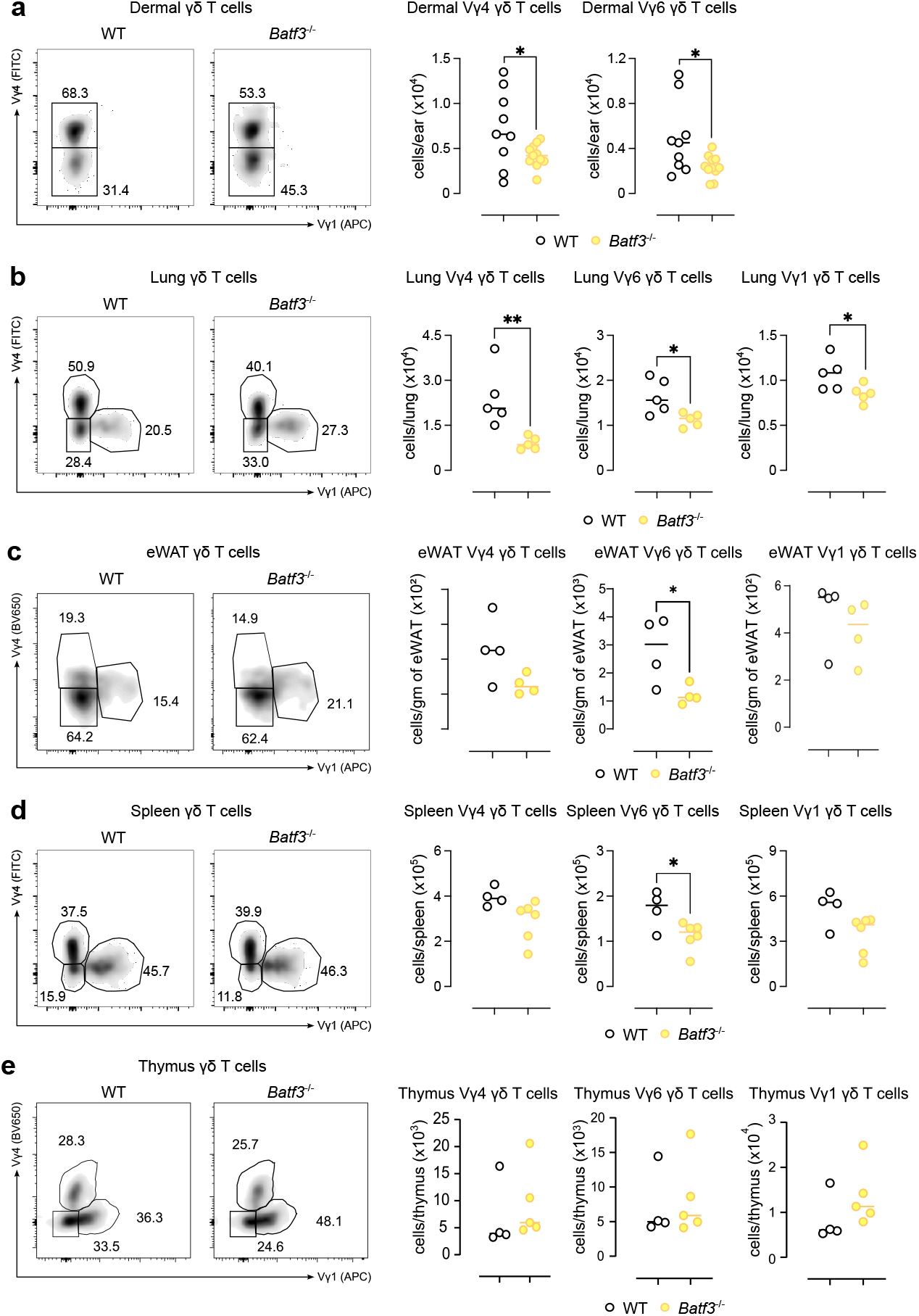
Distribution of γδ T cell subsets in skin, lung, eWAT, and thymus in WT and *Batf3*-deficient mice. Representative flow cytometry plots of γδ T cells subsets (Vγ4, Vγ6 and Vγ1) (left panels) and individual data showing the number of Vγ4, Vγ6 and Vγ1 γδ T cells (right panels) in the skin **(a)**, lung **(b)**, eWAT **(c)**, spleen **(d)** and thymus **(e). (a-e)** Data are presented as median (dots represent individual data points) and show a representative experiment. *, p <0.05; **, p < 0.01.

**Extended Data Figure 2.**
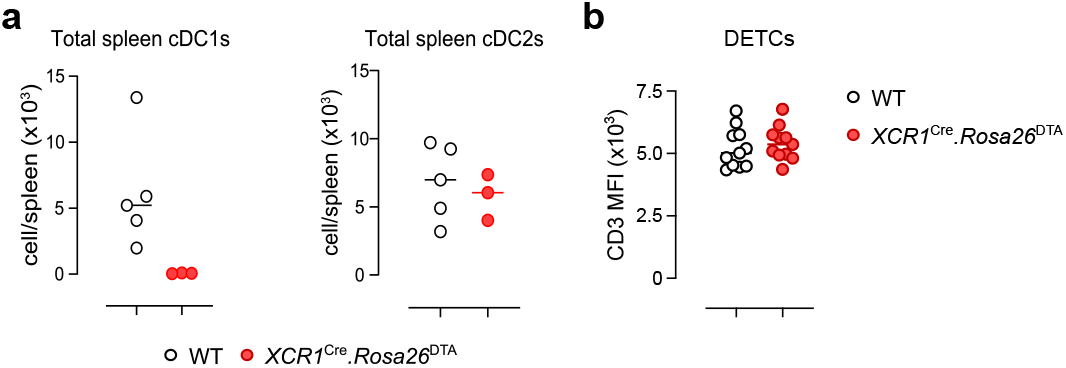
cDC1 cells are specifically depleted in XCR1-DTA^+^mice. **(a)** Analysis of percentages of cDC1s and cDC2s cells in the spleen of WT and XCR1-DTA+mice. **(b)** CD3 Median fluorescence intensity (MFI) in DETCs from WT and XCR1-DTA+mice. **(a, b)** Data are presented as median (dots represent individual data points) and show a pool of three experiments. **, p < 0.01.

**Extended Data Figure 3.**
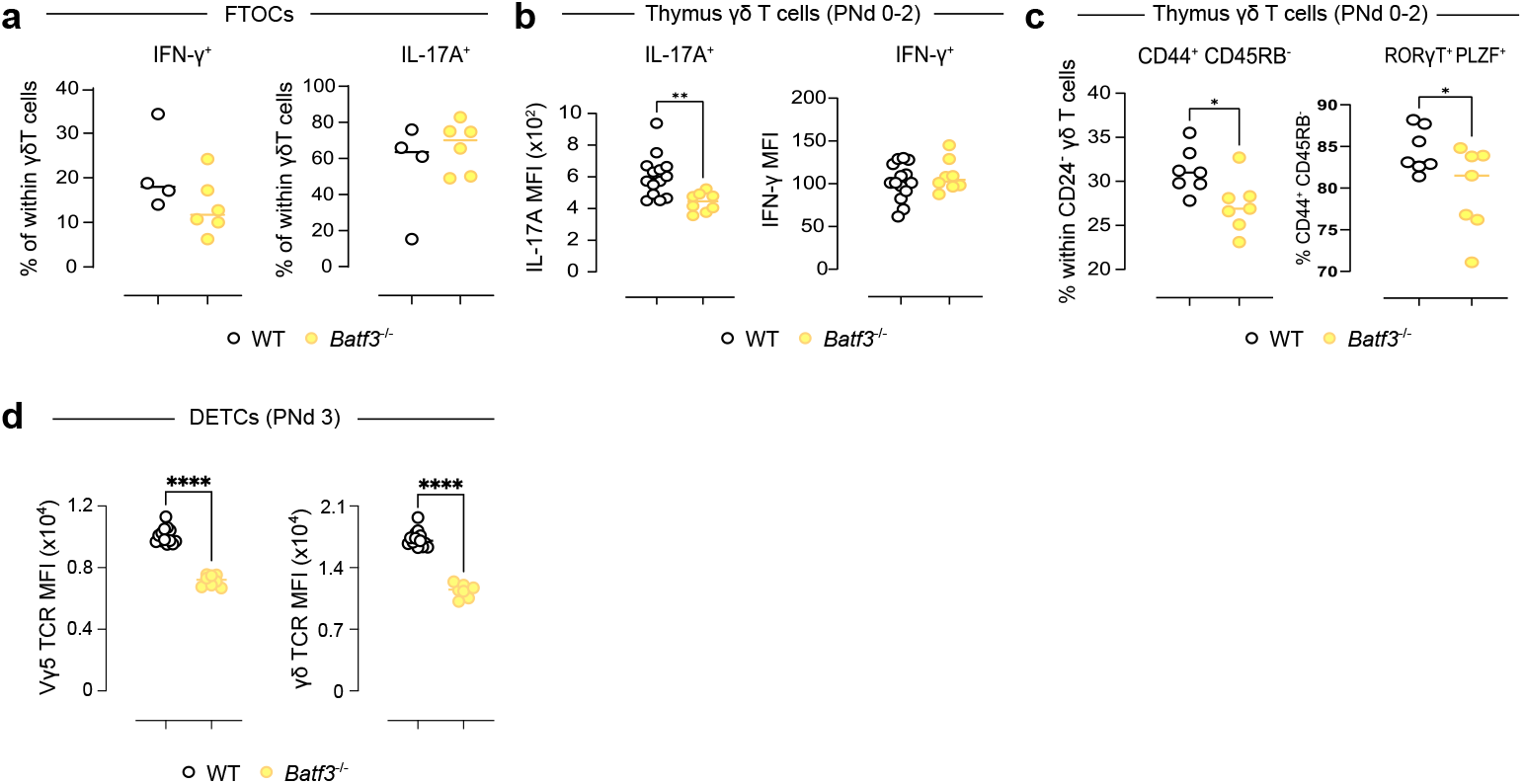
BATF3 deficiency does not affect γδ T cell development in the embryonic thymus. **(a)** IL17-A and IFN-γ production by embryionic γδ T cells in the HD-FTOCs of WT and *Batf3*-deficient mice. **(b)** Percentages of CD44^hi^CD45RB− γδ T cell subset and RORγt+PLZF+cells within the CD44^hi^CD45RB− γδ T cell subset in the neonatal thymus comparing WT and *Batf3*^−/−^mice. **(c)** MFI expression of IFN-γ and IL-17A in CD24^-^ γδ T cells of the neonatal thymus comparing WT and *Batf3*-deficient mice. **(d)** Vγ5 (left panel) and γδTCR (right panel) Median fluorescence intensity (MFI) in DETCs from WT and and *Batf3*^−/−^ mice. **(a-d)** Data are presented as median (dots represent individual data points) and a representative experiment. *, p <0.05; **, p < 0.01; ****, p< 0.0001.

**Extended Data Figure 4.**
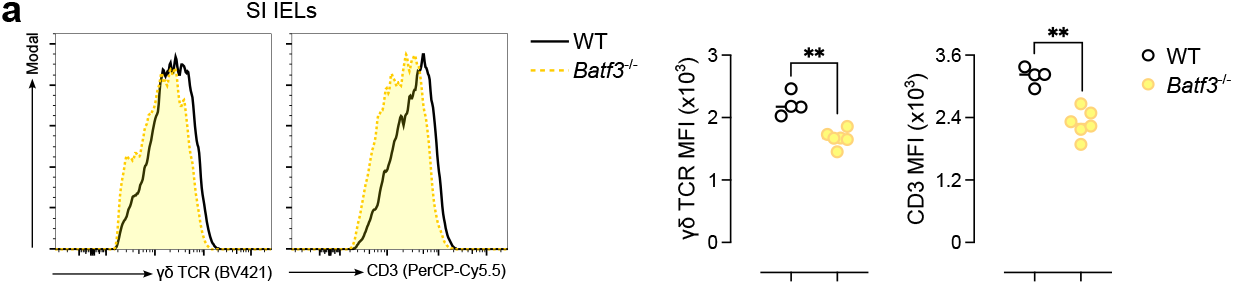
Impaired TCR expression in γδ T cells from *Batf3*-deficient mice. **(a)** Representative histograms (left panels) and summary plots (right panels) of TCR complex molecules median fluorescence intensity (MFI) in intraepithelial γδ T cells (SI IELs) from WT and *Batf3*^−*/*−^ mice. Data are presented as median (dots represent individual data points) and is representative of at least 2 independent experiments. Normality of the samples was assessed, and statistical analysis was then performed accordingly. **, *p* < 0.01

**Extended Data Figure 5.**
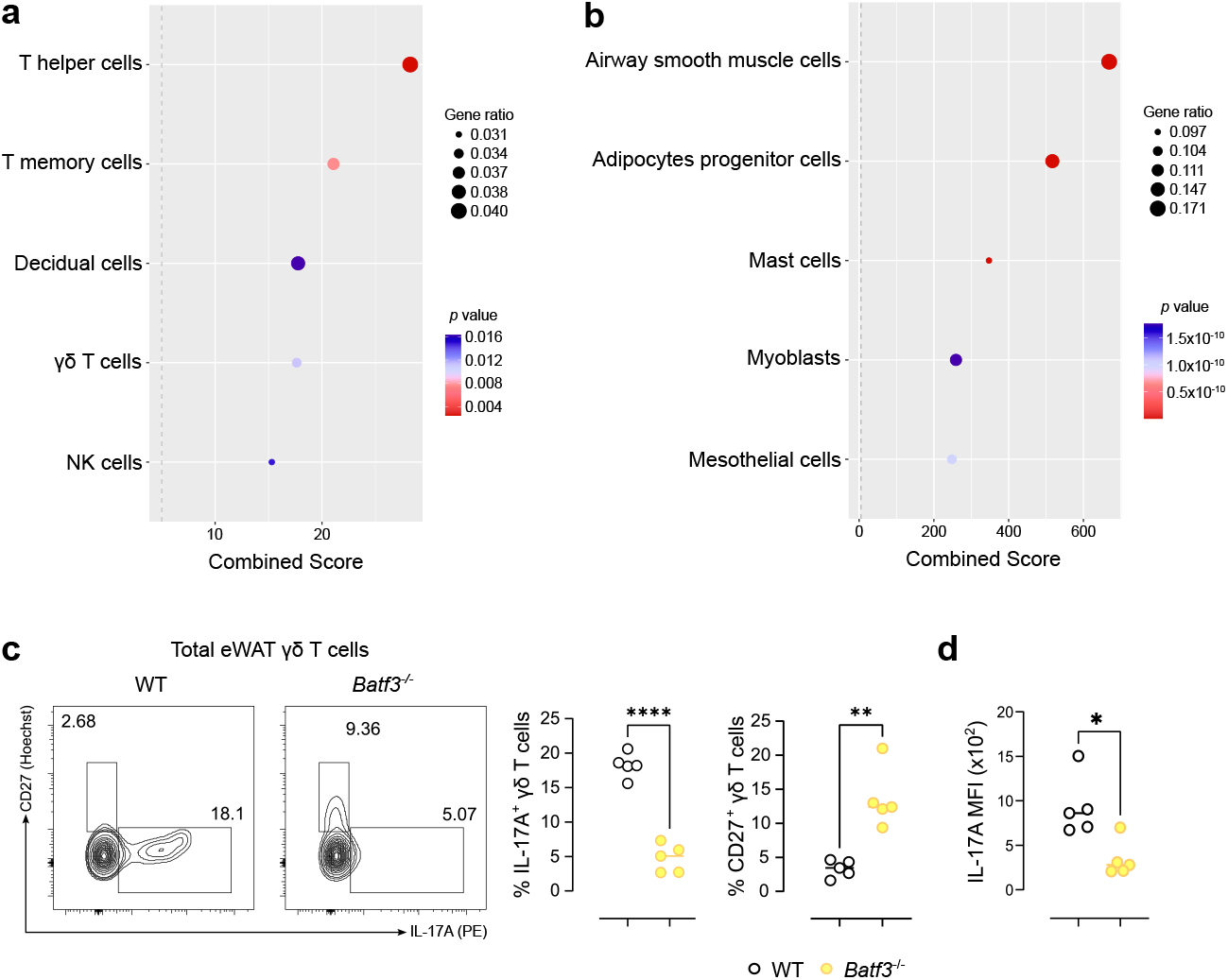
BATF3 regulates γδ17 T cell identity. **(a, b)** Cell identities extracted from the PanglaoDB database for scRNAseq studies using significantly upregulated genes from **(a)** WT dermal γδ T cells and **(b)** *Batf3*^−*/*−^ dermal γδ T cells. **(c)** Representative flow cytometry plots (left panels) and summary plots (right panels) depicting IL-17A production and CD27 expression by γδ T cells in the eWAT after 4h of ex-vivo stimulation comparing WT and *Batf3*^−*/*−^ mice. Data are presented as median (dots represent individual data points). **(d)** Median fluorescence intensity (MFI) expression of de IL-17A in γδ T cells of the eWAT comparing WT and *Batf3*-deficient mice. **(c–d)** Data are presented as median; each dot represents an individual mouse. Data representative of at least 2 independent experiments. Normality of the samples was assessed, and statistical analysis was then performed accordingly. *, p <0.05; **, *p* < 0.01; ****, *p* < 0.0001.

**Extended Data Table 1.**
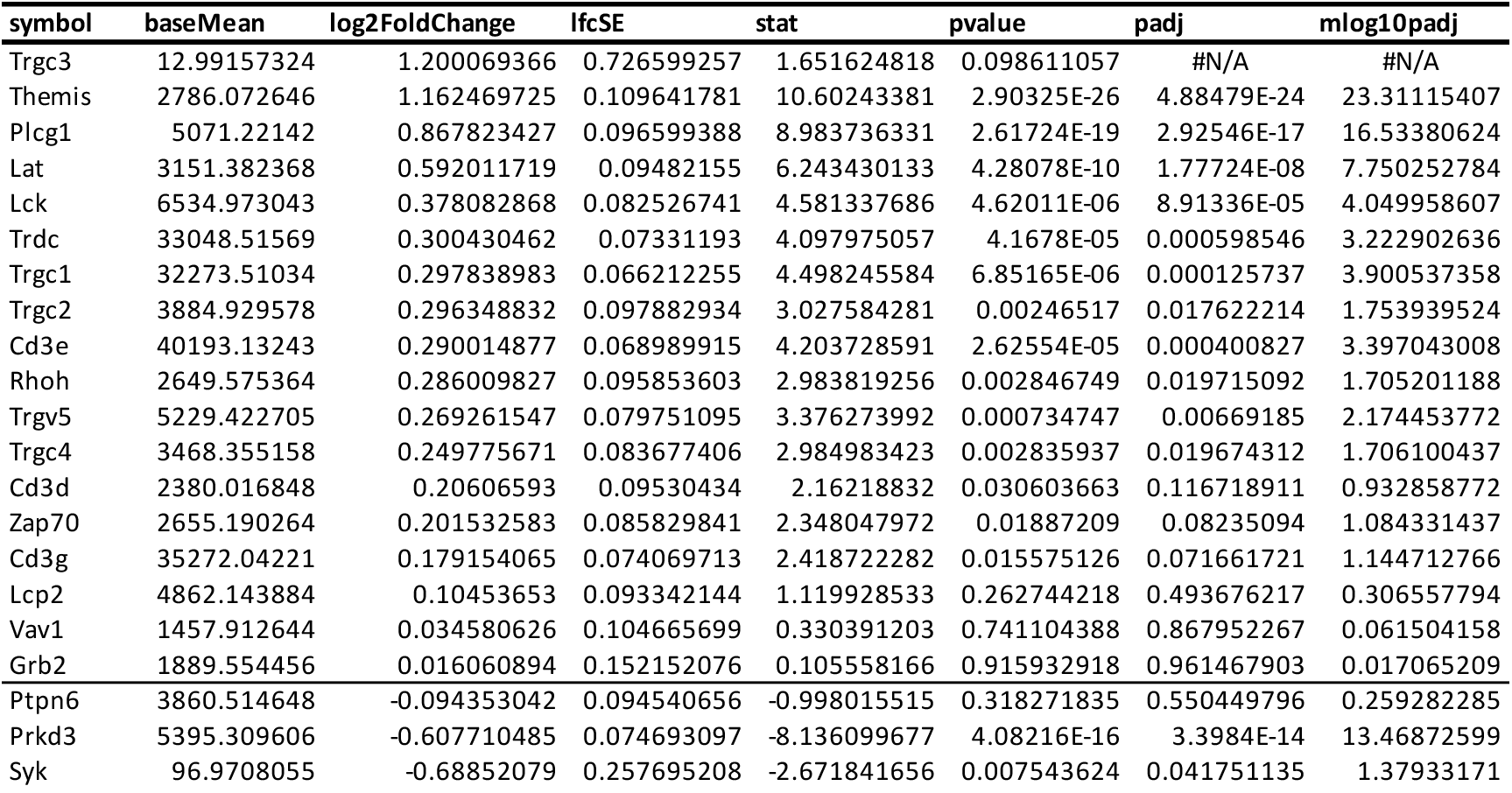
TCR complex genes (WT vs KO)

| symbol | baseMean | log2FoldChange | lfcSE | stat | pvalue | padj | mlog10padj |
| --- | --- | --- | --- | --- | --- | --- | --- |
| Trgc3 | 12.99157324 | 1.200069366 | 0.726599257 | 1.651624818 | 0.098611057 | #N/A | #N/A |
| Themis | 2786.072646 | 1.162469725 | 0.109641781 | 10.60243381 | 2.90325E-26 | 4.88479E-24 | 23.31115407 |
| Plcg1 | 5071.22142 | 0.867823427 | 0.096599388 | 8.983736331 | 2.61724E-19 | 2.92546E-17 | 16.53380624 |
| Lat | 3151.382368 | 0.592011719 | 0.09482155 | 6.243430133 | 4.28078E-10 | 1.77724E-08 | 7.750252784 |
| Lck | 6534.973043 | 0.378082868 | 0.082526741 | 4.581337686 | 4.62011E-06 | 8.91336E-05 | 4.049958607 |
| Trdc | 33048.51569 | 0.300430462 | 0.07331193 | 4.097975057 | 4.1678E-05 | 0.000598546 | 3.222902636 |
| Trgc1 | 32273.51034 | 0.297838983 | 0.066212255 | 4.498245584 | 6.85165E-06 | 0.000125737 | 3.900537358 |
| Trgc2 | 3884.929578 | 0.296348832 | 0.097882934 | 3.027584281 | 0.00246517 | 0.017622214 | 1.753939524 |
| Cd3e | 40193.13243 | 0.290014877 | 0.068989915 | 4.203728591 | 2.62554E-05 | 0.000400827 | 3.397043008 |
| Rhoh | 2649.575364 | 0.286009827 | 0.095853603 | 2.983819256 | 0.002846749 | 0.019715092 | 1.705201188 |
| Trgv5 | 5229.422705 | 0.269261547 | 0.079751095 | 3.376273992 | 0.000734747 | 0.00669185 | 2.174453772 |
| Trgc4 | 3468.355158 | 0.249775671 | 0.083677406 | 2.984983423 | 0.002835937 | 0.019674312 | 1.706100437 |
| Cd3d | 2380.016848 | 0.20606593 | 0.09530434 | 2.16218832 | 0.030603663 | 0.116718911 | 0.932858772 |
| Zap70 | 2655.190264 | 0.201532583 | 0.085829841 | 2.348047972 | 0.01887209 | 0.08235094 | 1.084331437 |
| Cd3g | 35272.04221 | 0.179154065 | 0.074069713 | 2.418722282 | 0.015575126 | 0.071661721 | 1.144712766 |
| Lcp2 | 4862.143884 | 0.10453653 | 0.093342144 | 1.119928533 | 0.262744218 | 0.493676217 | 0.306557794 |
| Vav1 | 1457.912644 | 0.034580626 | 0.104665699 | 0.330391203 | 0.741104388 | 0.867952267 | 0.061504158 |
| Grb2 | 1889.554456 | 0.016060894 | 0.152152076 | 0.105558166 | 0.915932918 | 0.961467903 | 0.017065209 |
| Ptpn6 | 3860.514648 | -0.094353042 | 0.094540656 | -0.998015515 | 0.318271835 | 0.550449796 | 0.259282285 |
| Prkd3 | 5395.309606 | -0.607710485 | 0.074693097 | -8.136099677 | 4.08216E-16 | 3.3984E-14 | 13.46872599 |
| Syk | 96.9708055 | -0.68852079 | 0.257695208 | -2.671841656 | 0.007543624 | 0.041751135 | 1.37933171 |

**Extended Data Table 2.**
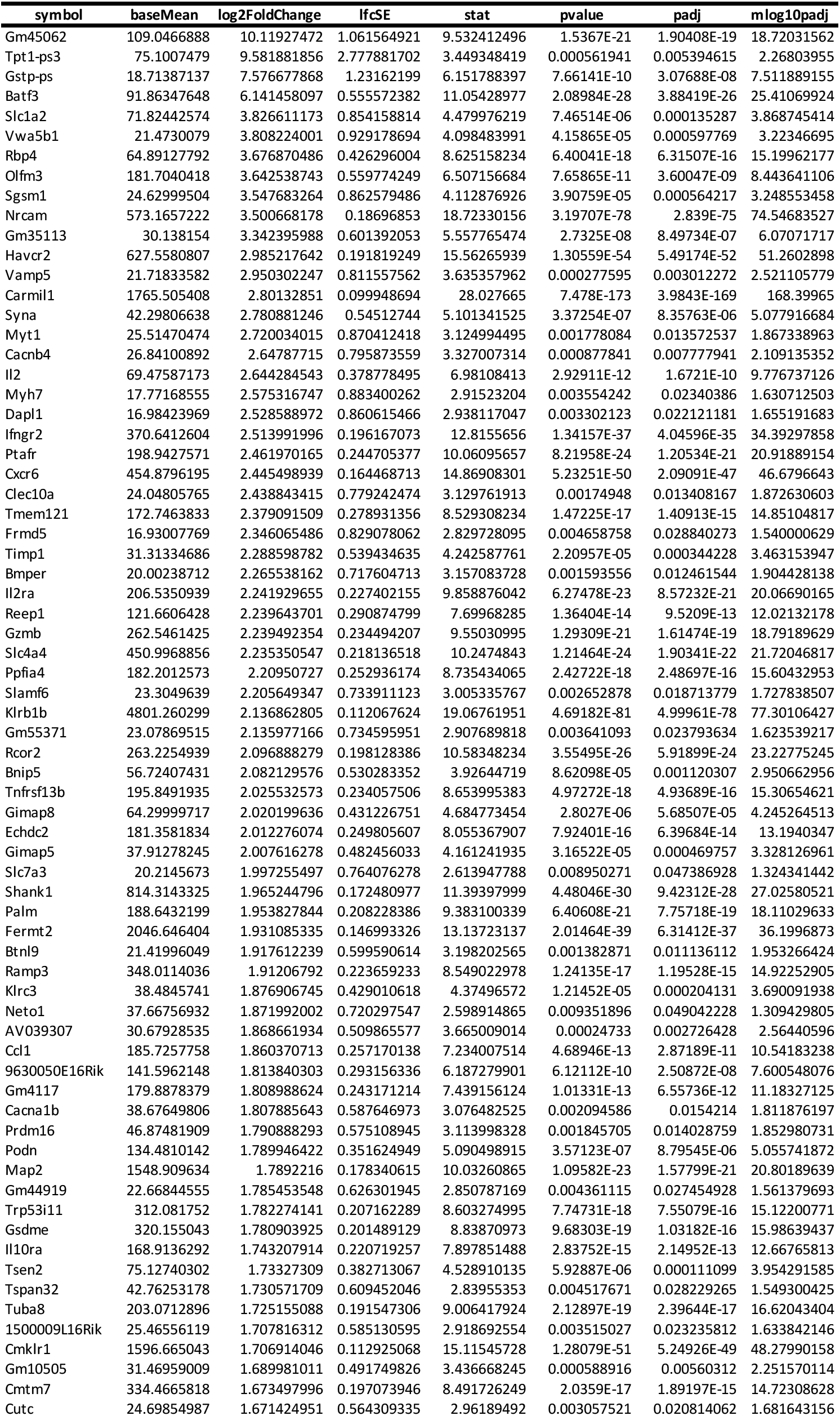
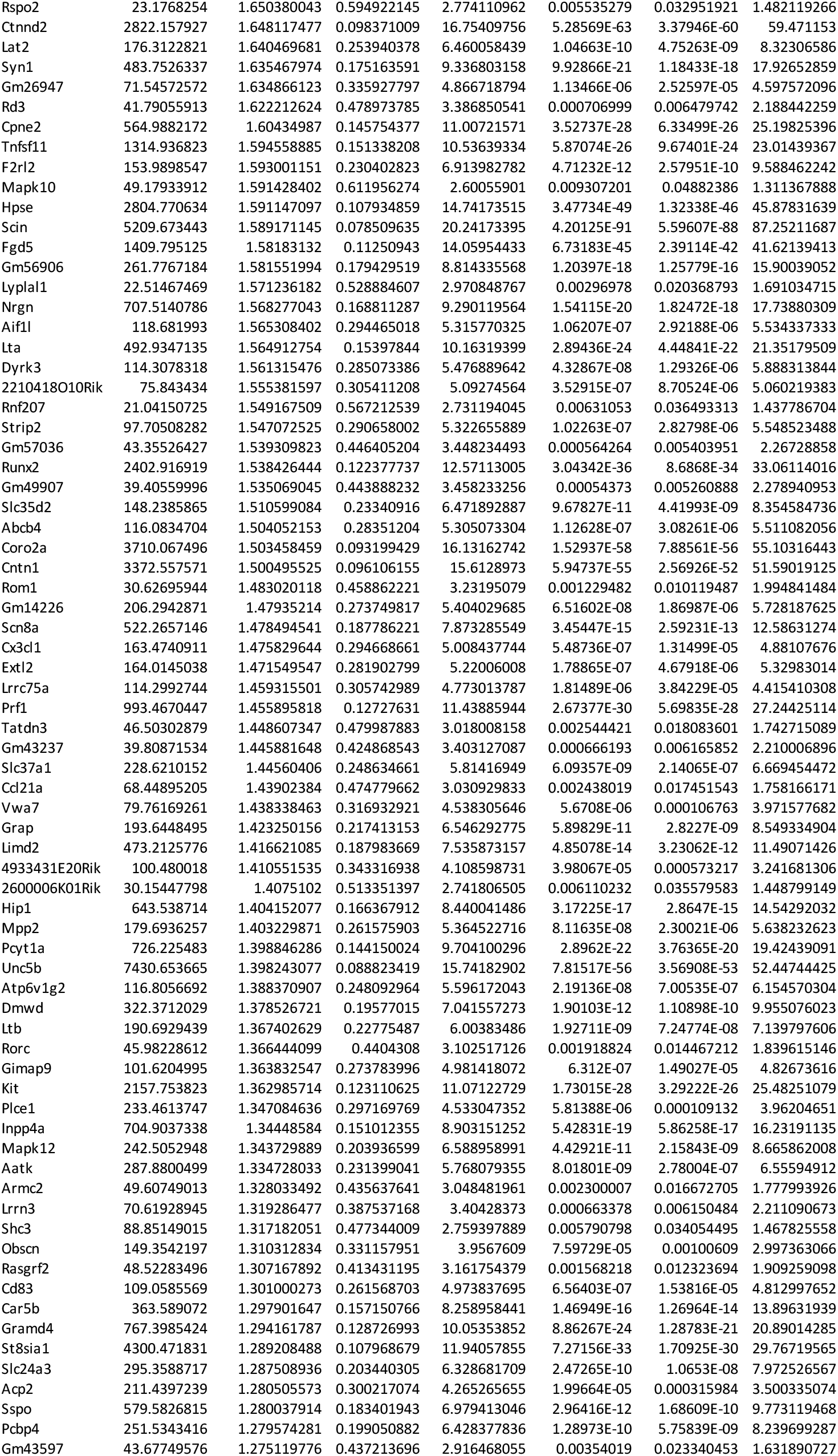
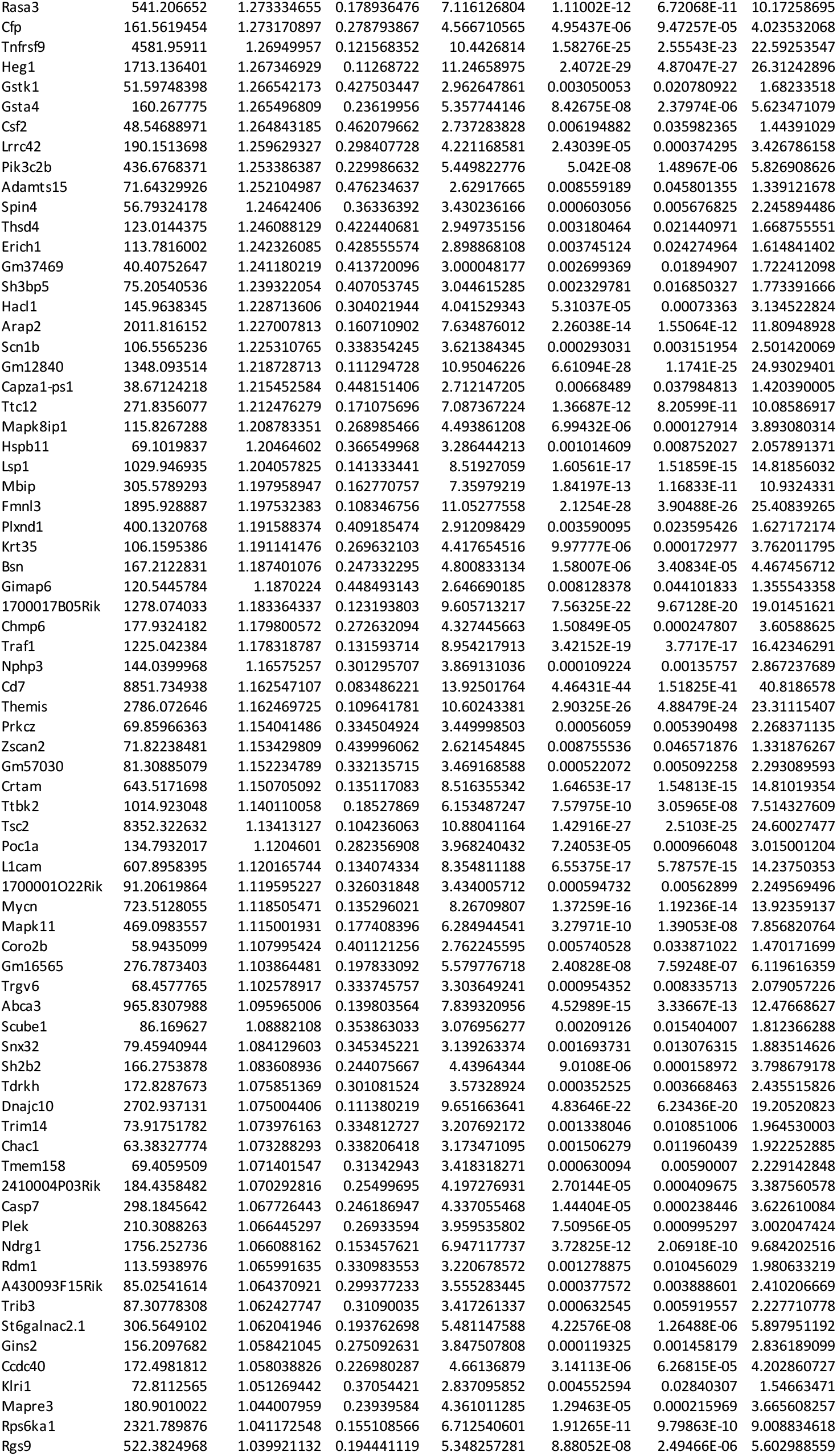
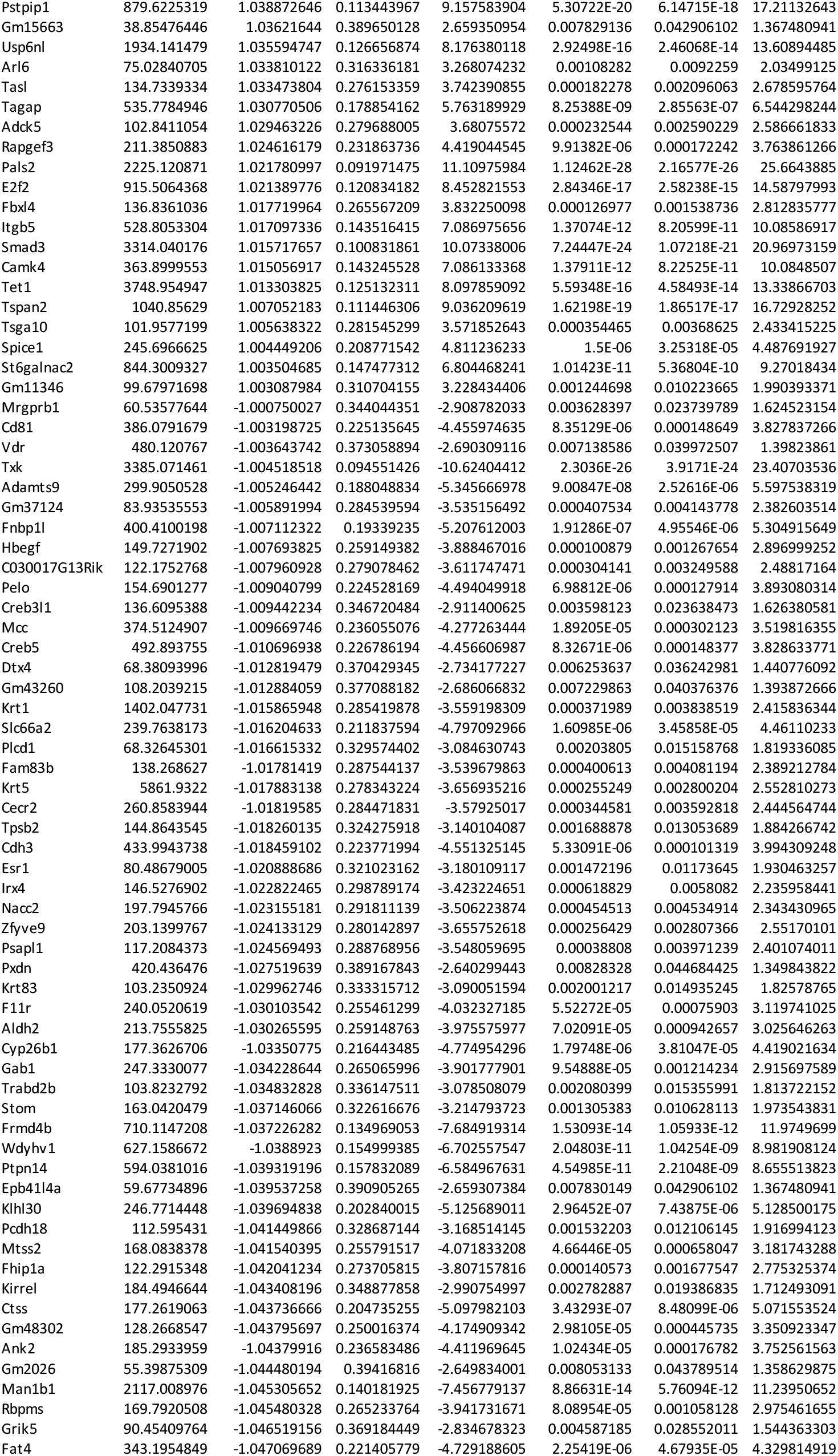
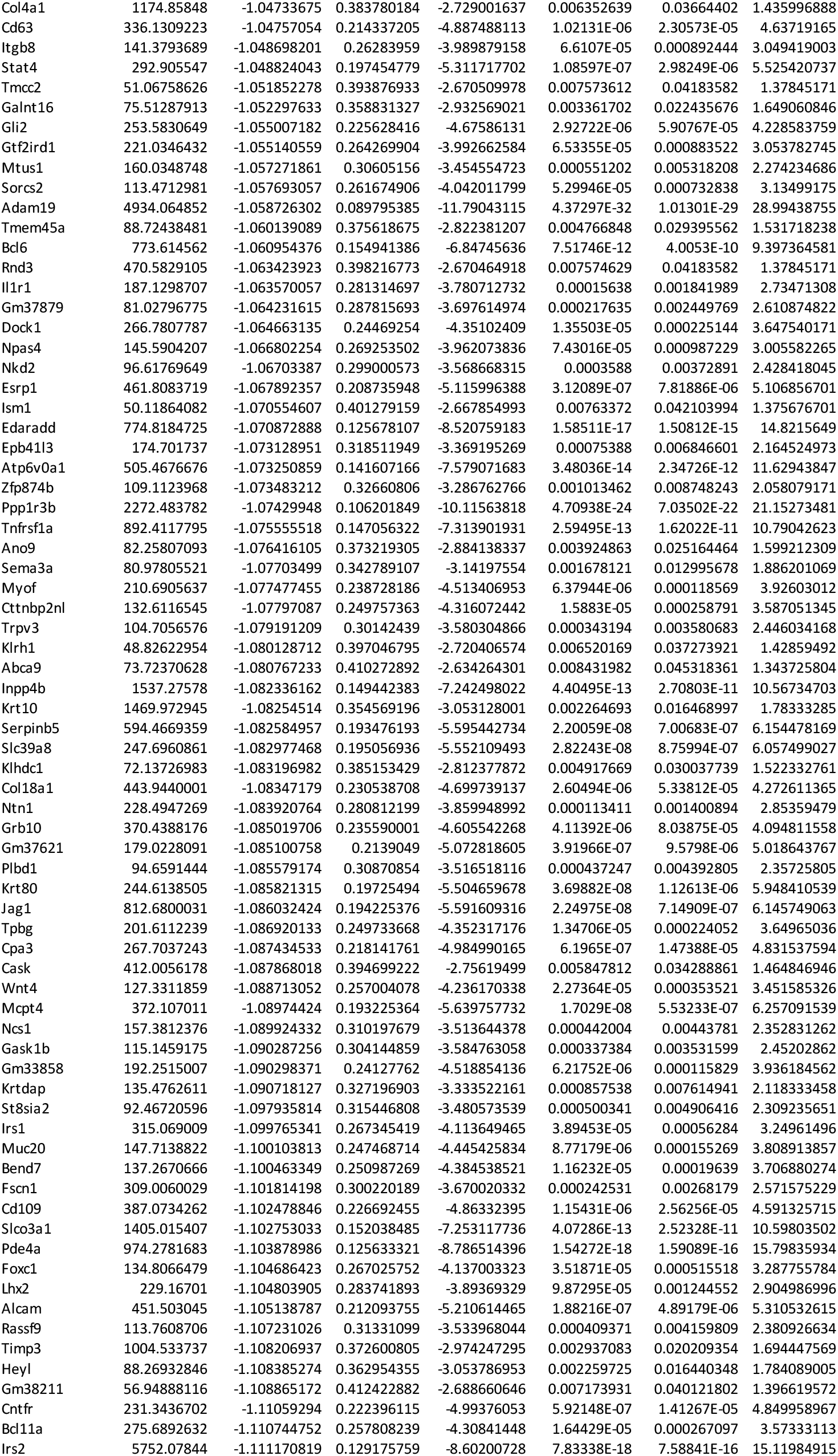
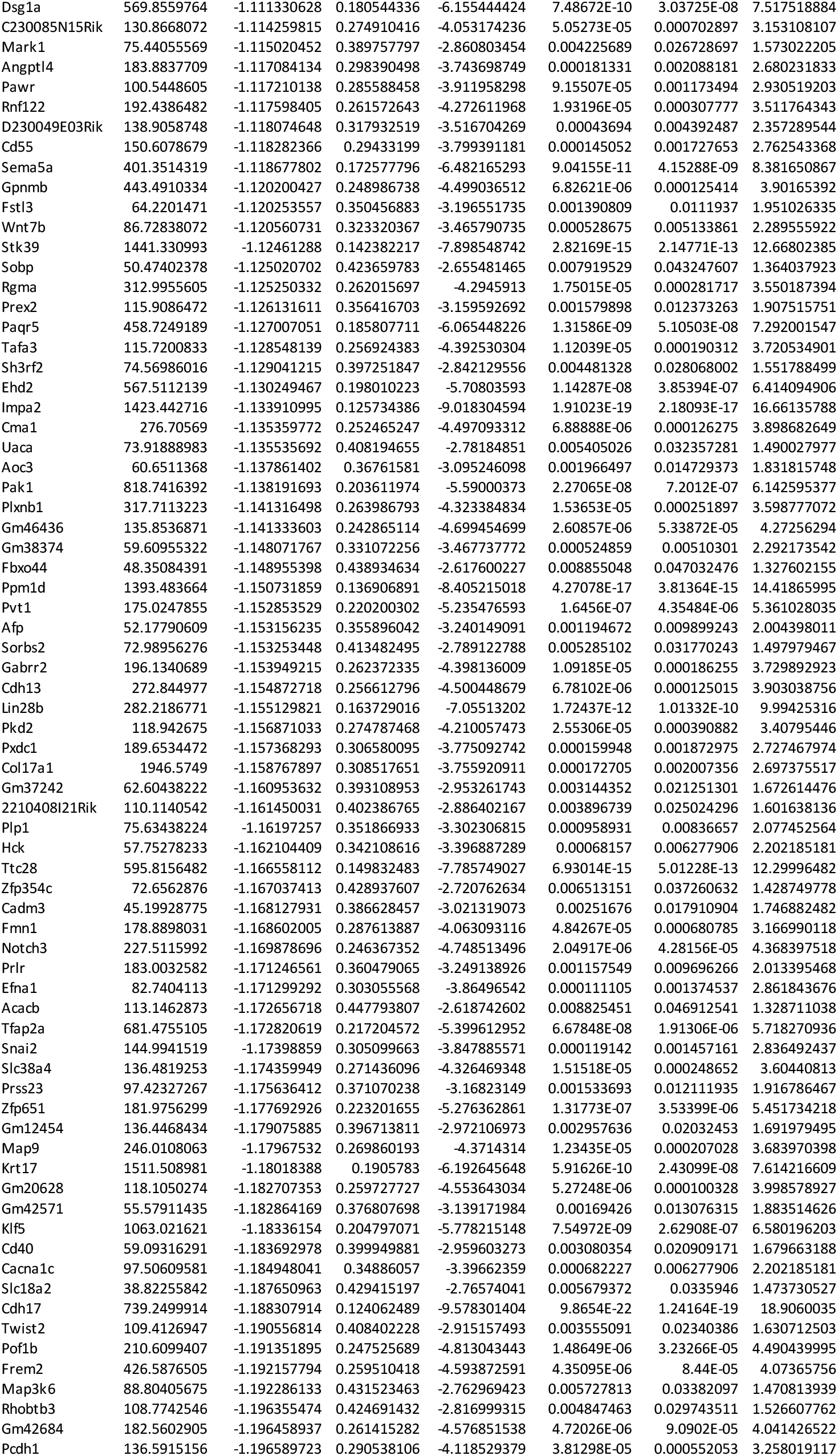
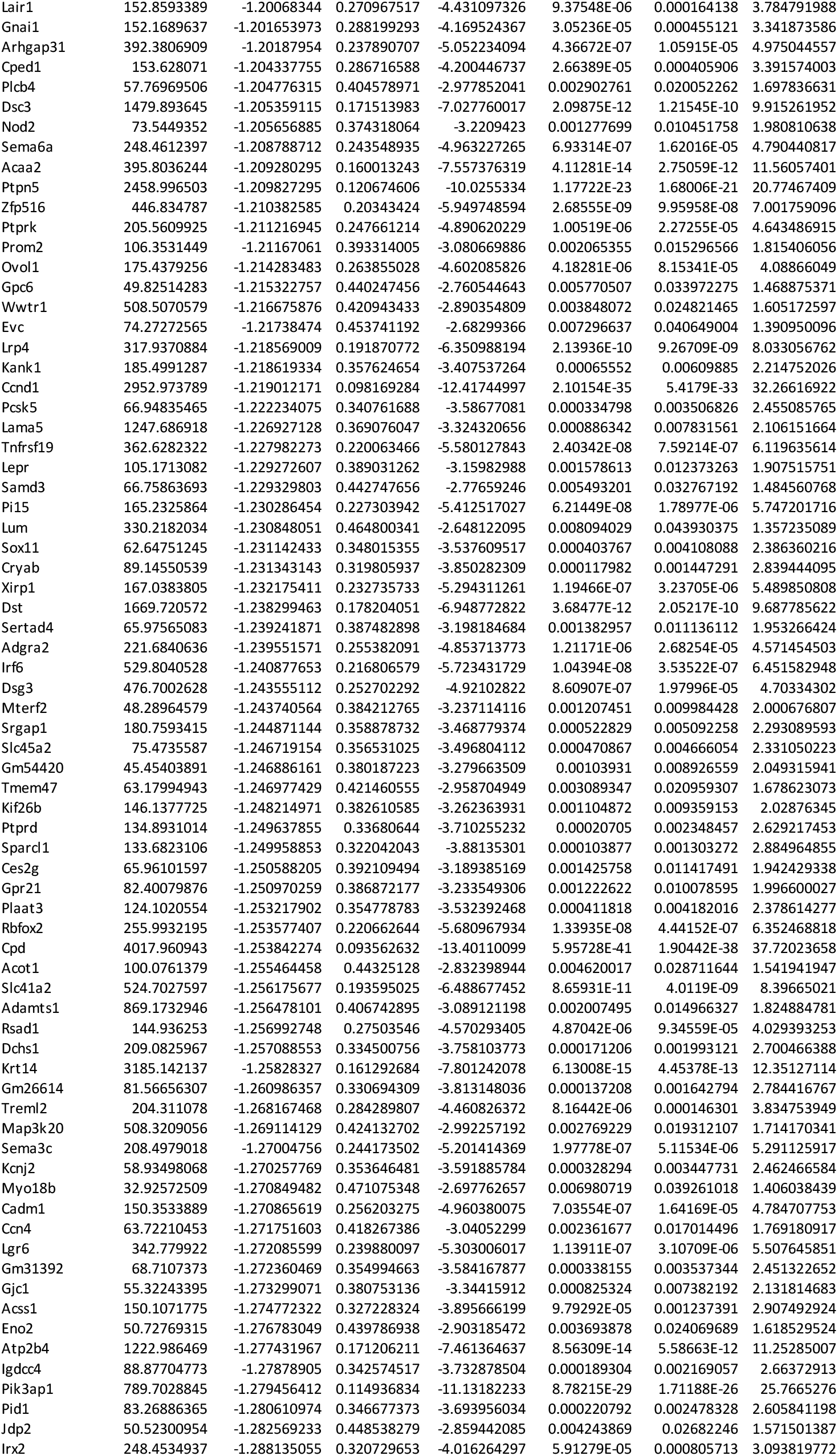
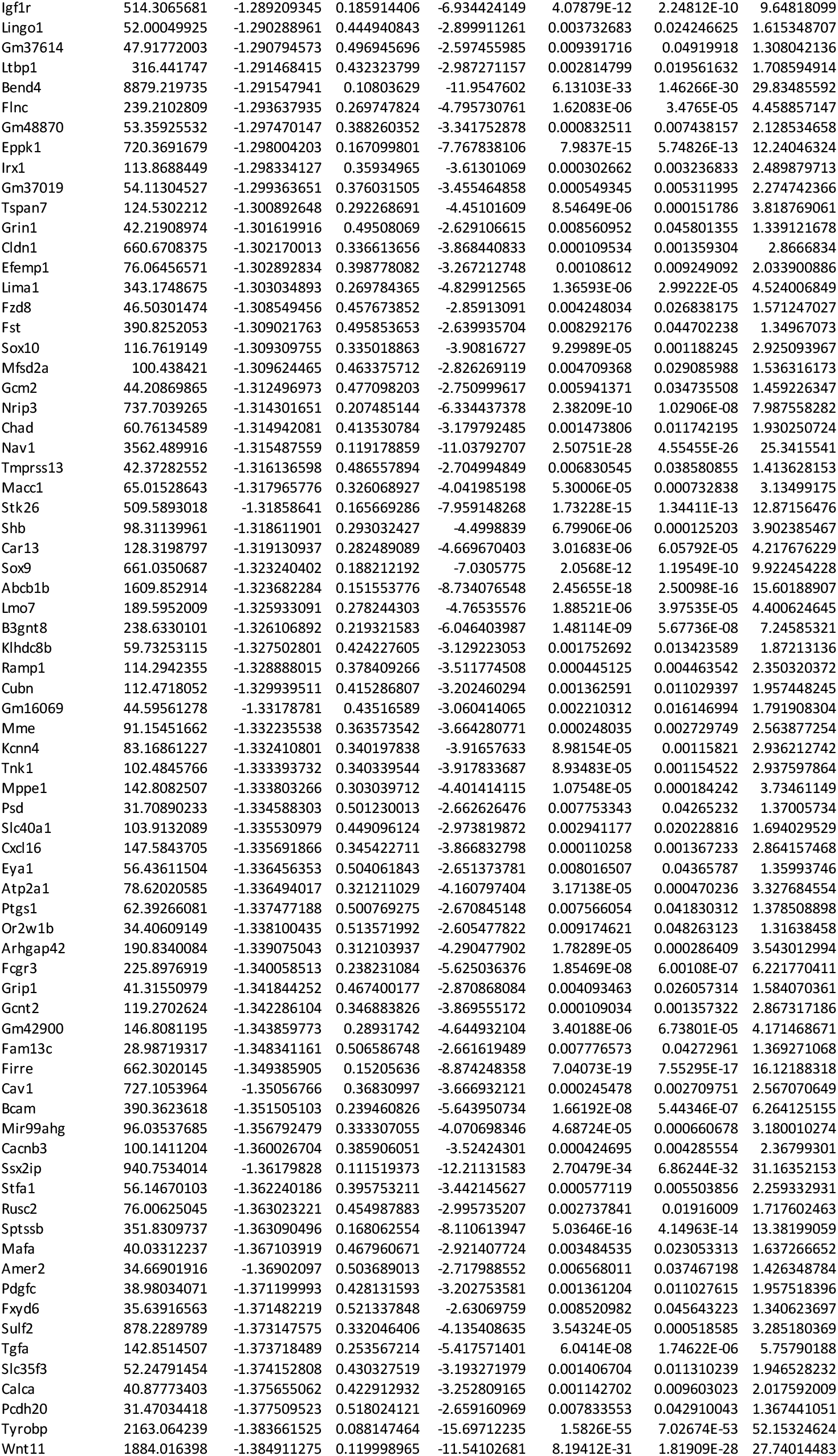
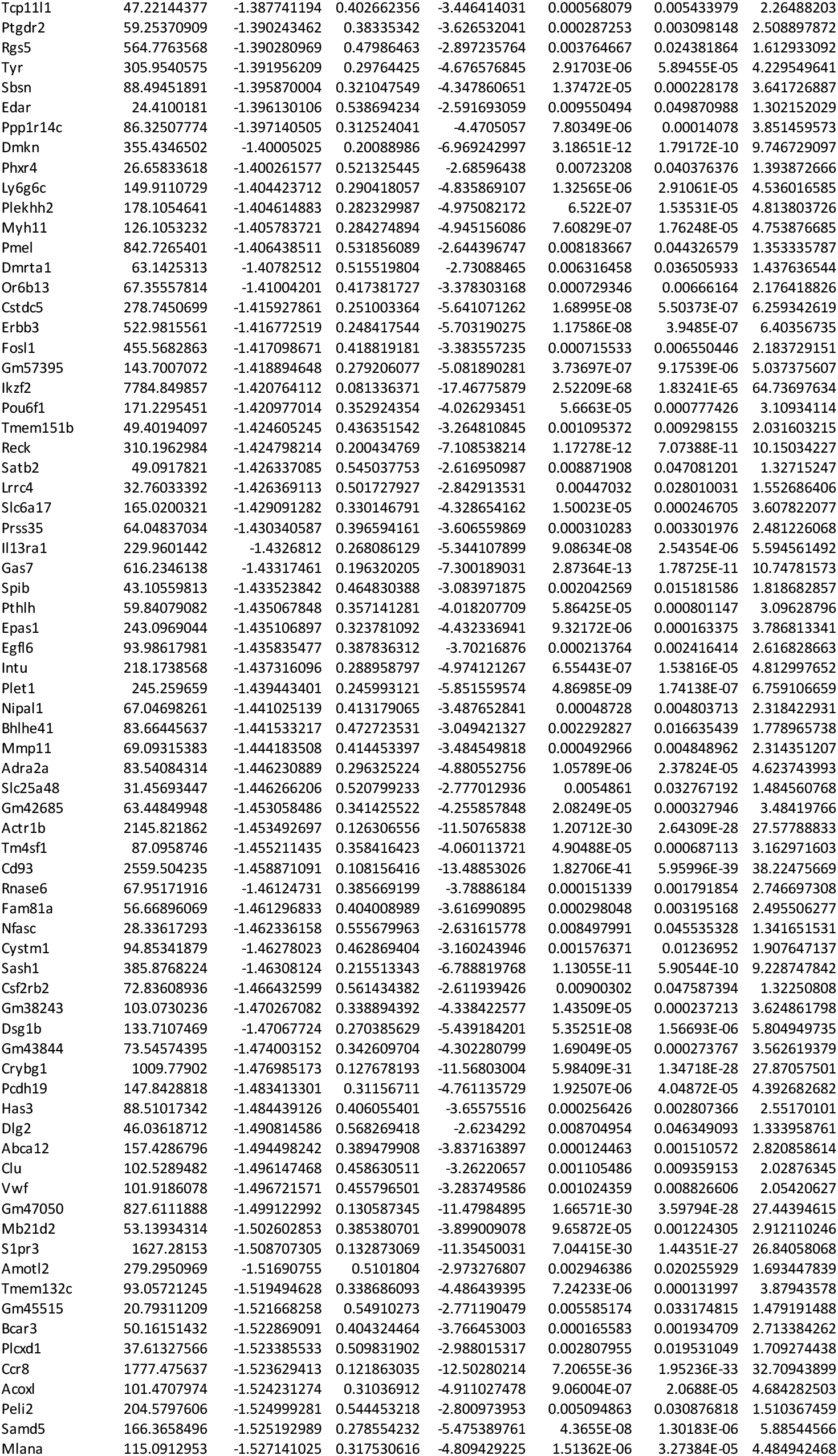
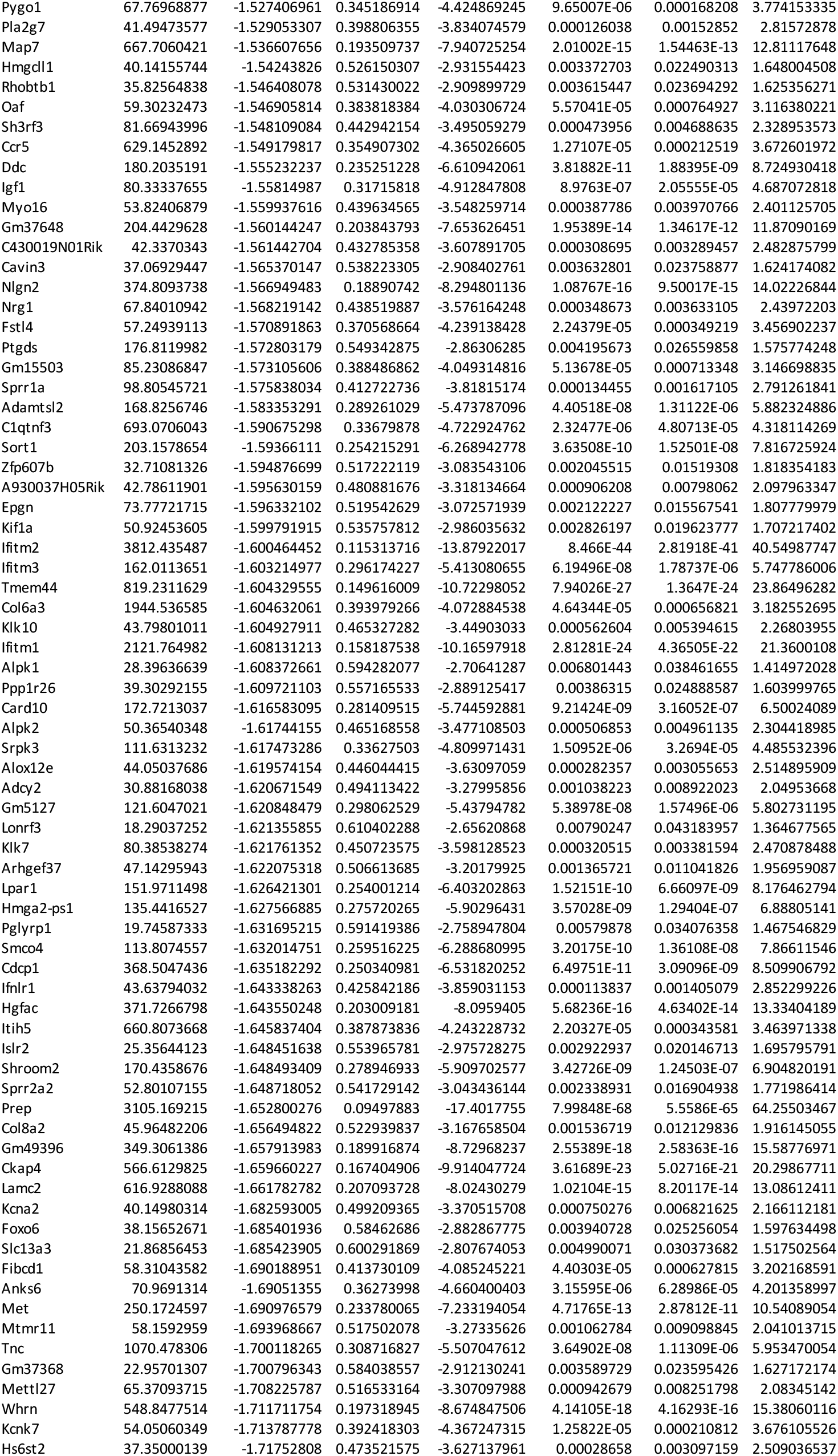
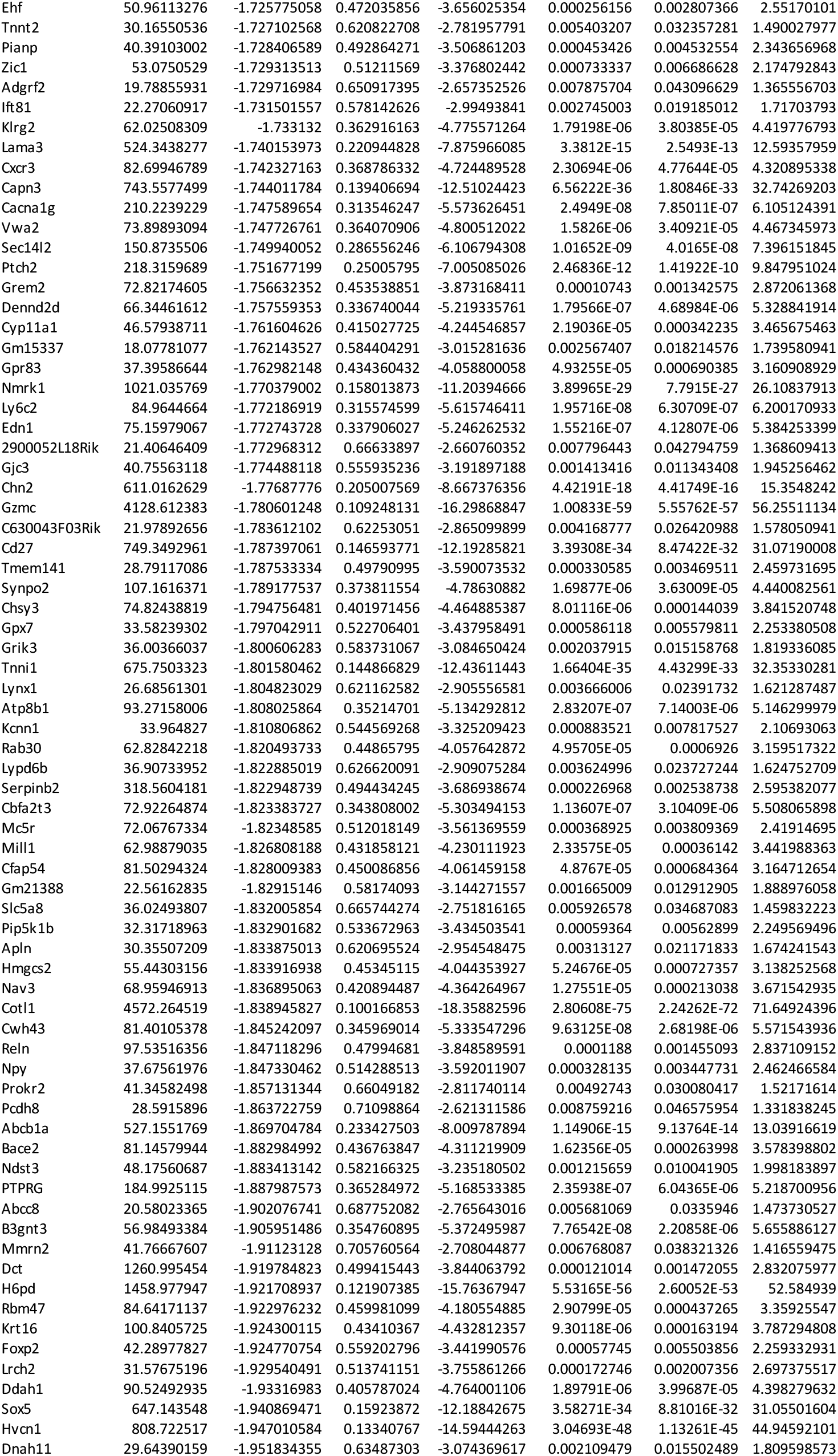
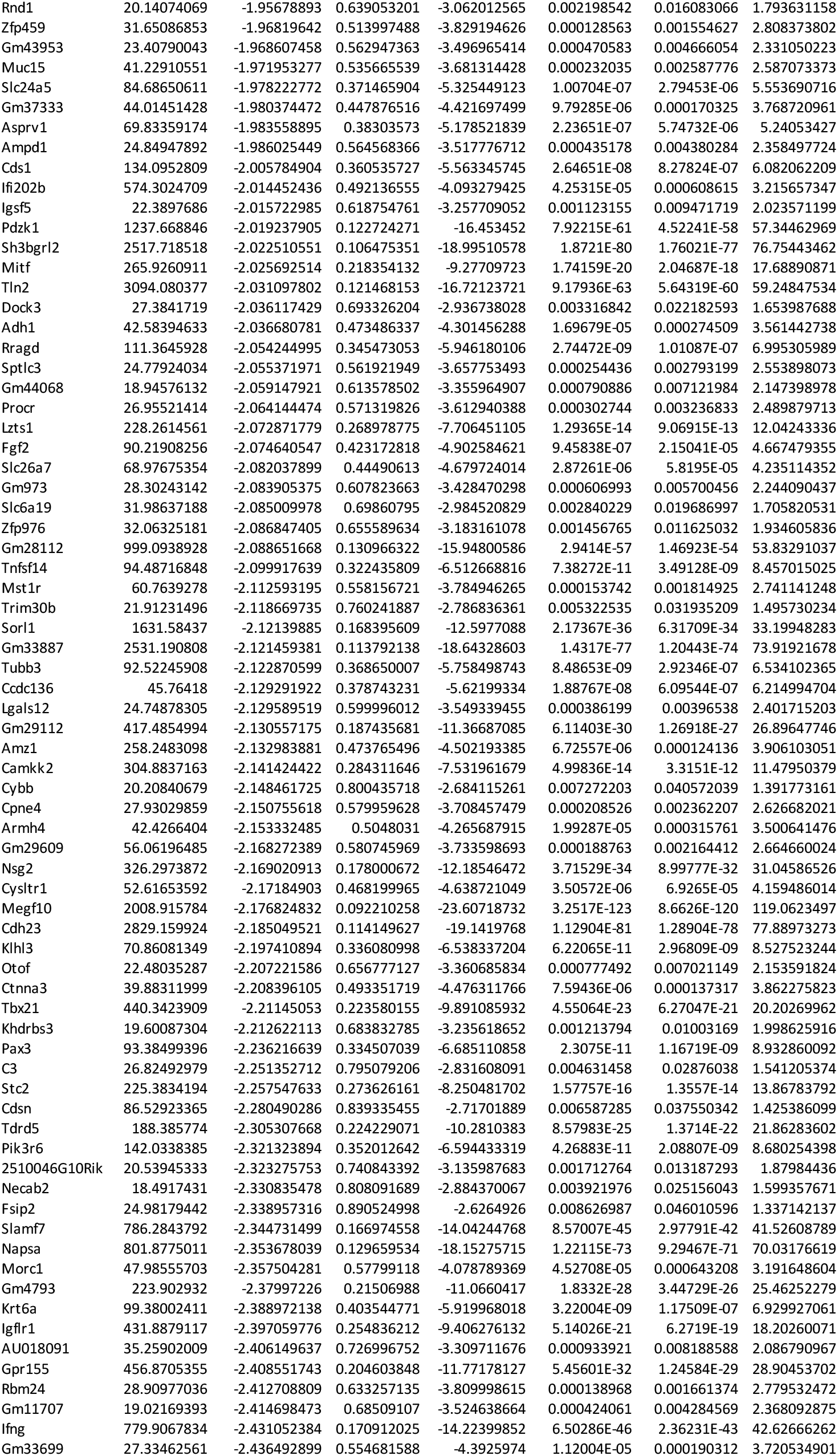
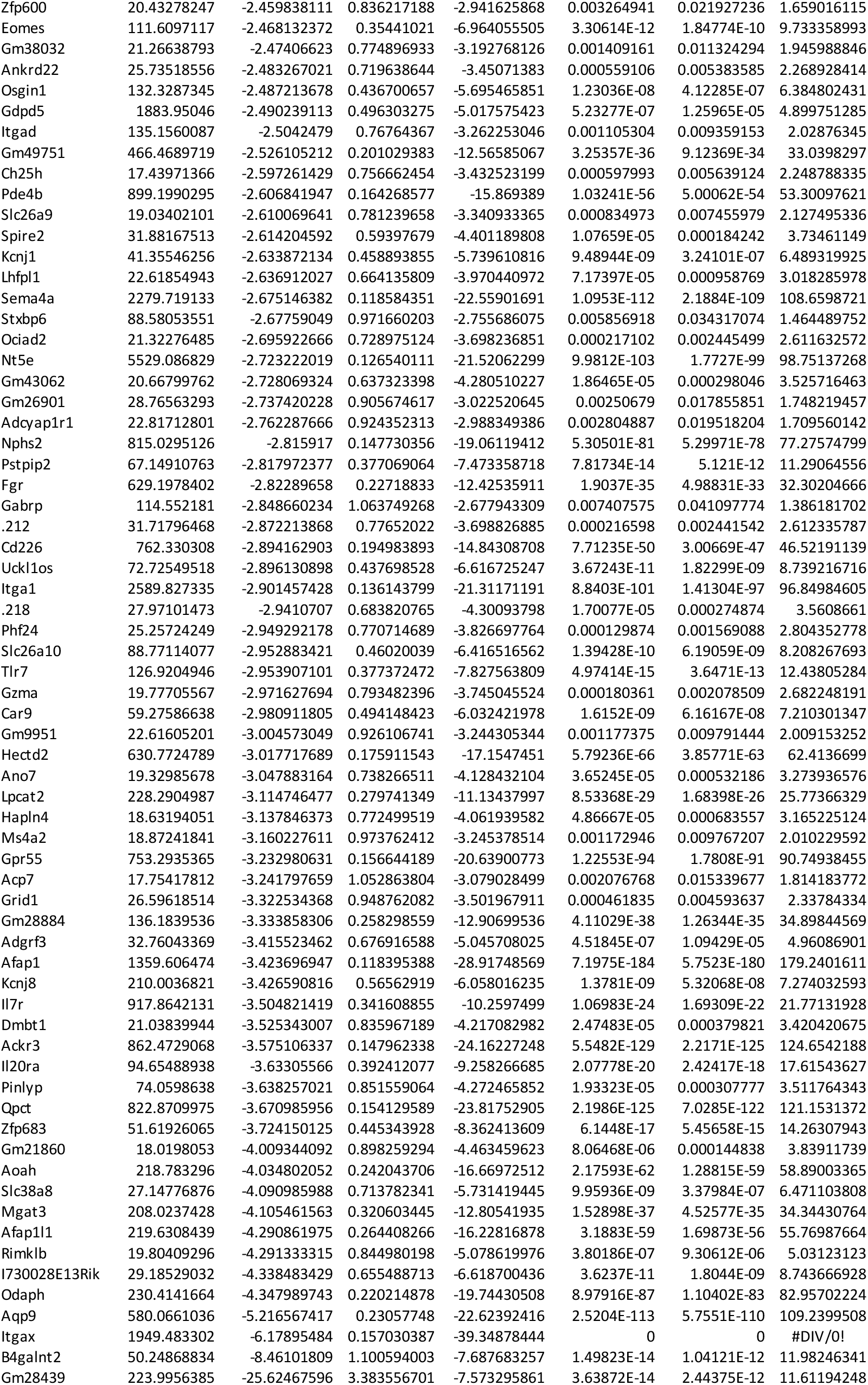
DEGs between WT and *Batf3* -/-DETCs.

**Extended Data Table 3.**
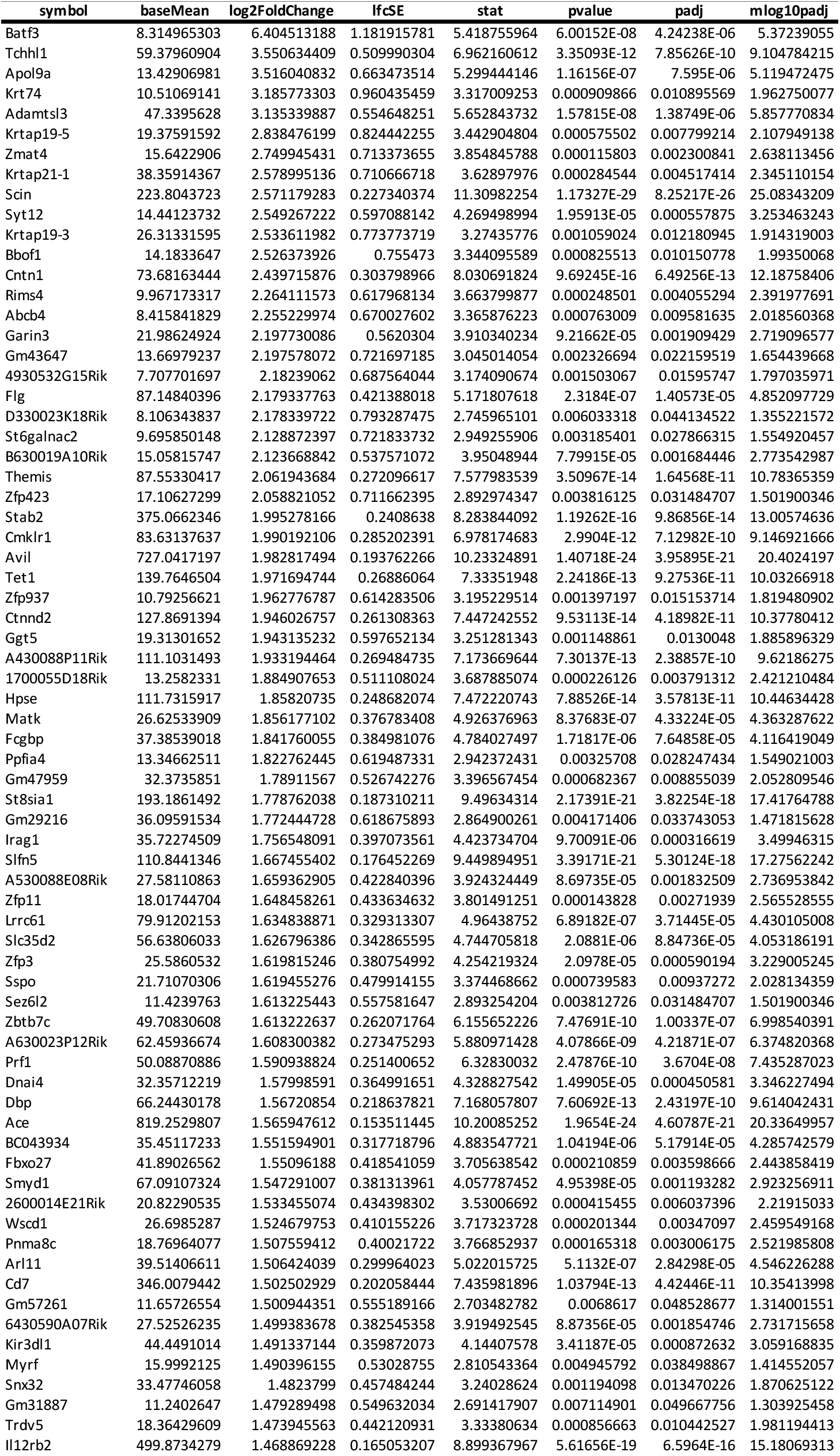
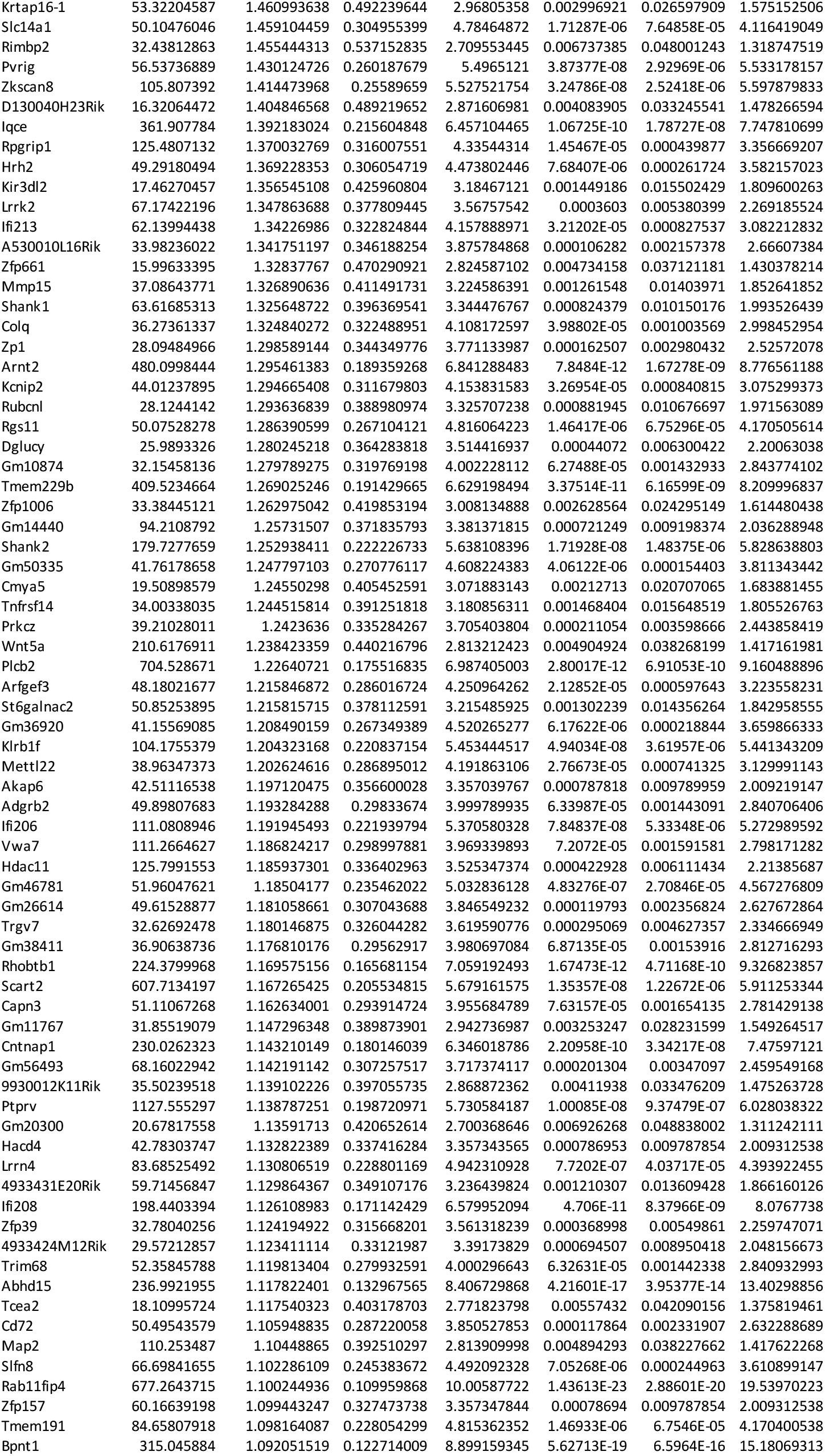
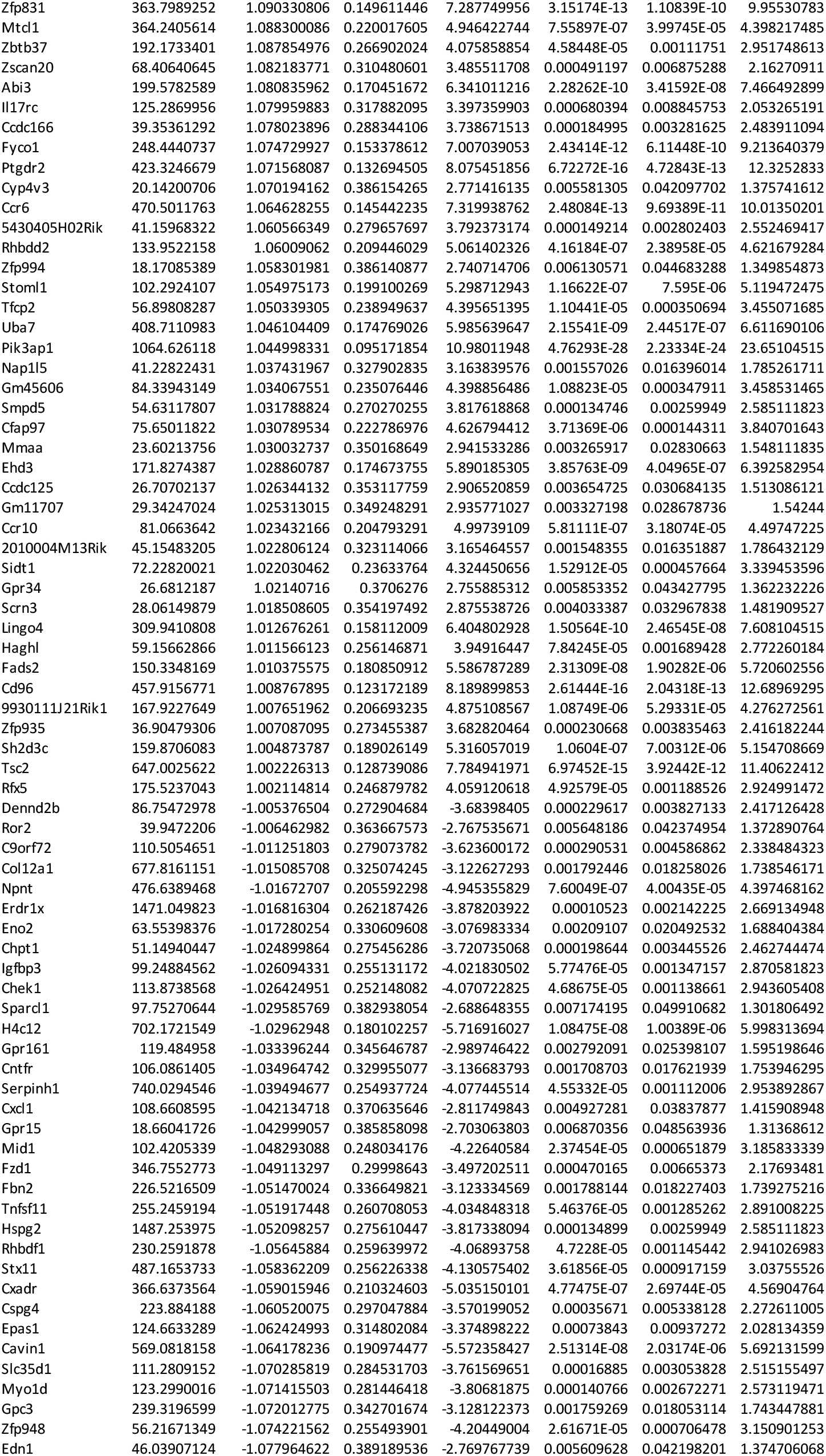
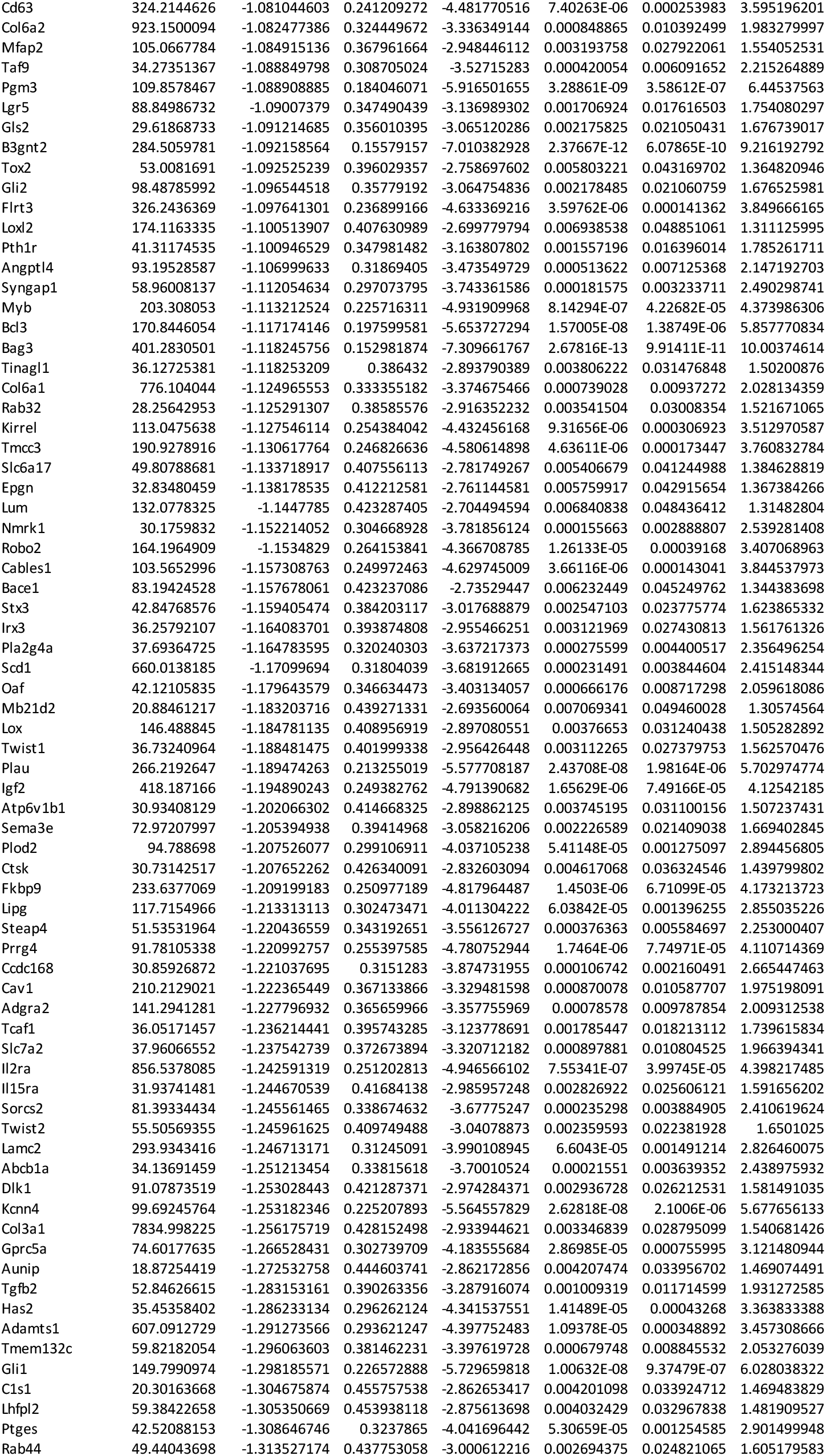
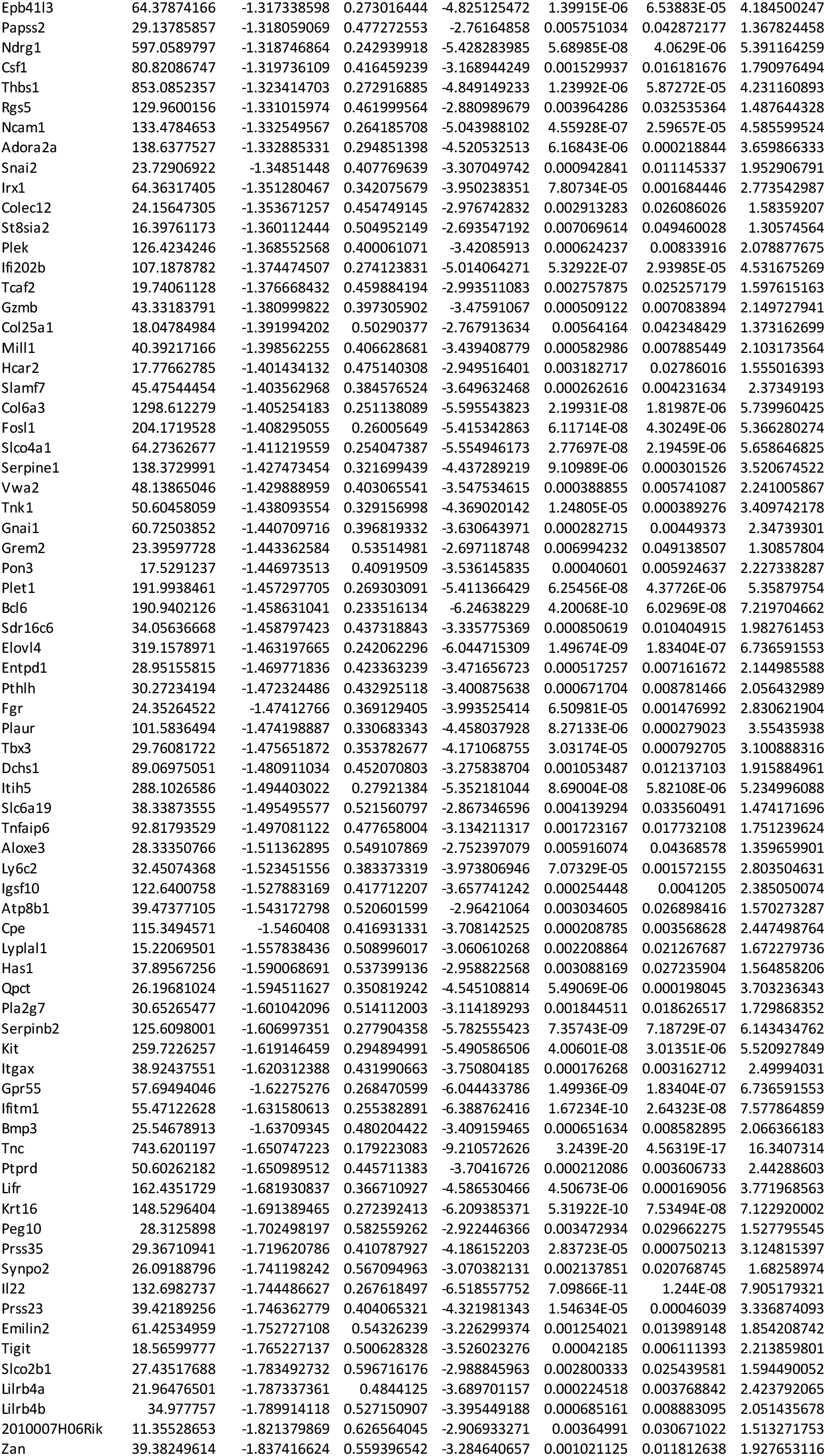
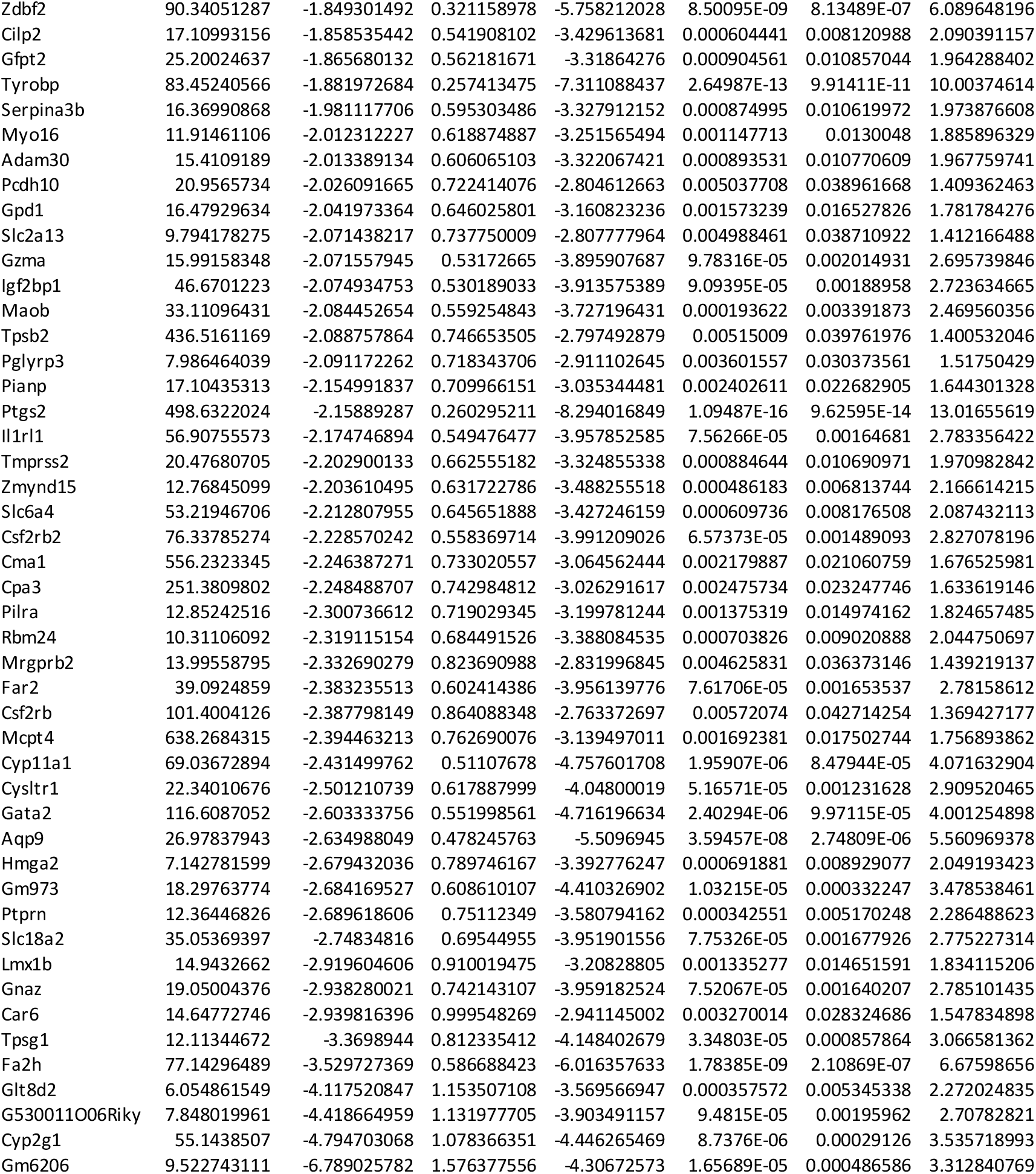
DEGs between WT and *Batf3* -/-dermal gd T cells.

**Extended Data Table 4.** γδ17 T cell signature genes (WT vs KO)

| symbol | baseMean | log2FoldChange | lfcSE | stat | pvalue | padj |
| --- | --- | --- | --- | --- | --- | --- |
| Il22 | 132.6982737 | -1.744486627 | 0.267618 | -6.518557752 | 7.1E-11 | 1.244E-08 |
| Il17a | 231.1708114 | -0.658246096 | 0.244006 | -2.697660913 | 0.006983 | 0.049113895 |
| Gpr183 | 620.9806588 | 0.043975157 | 0.168936 | 0.260306482 | 0.794627 | 0.910202883 |
| Il23r | 1240.666281 | 0.391656208 | 0.134776 | 2.905986049 | 0.003661 | 0.030708988 |
| Sox13 | 1480.137565 | 0.622317894 | 0.134873 | 4.614097822 | 3.95E-06 | 0.000150917 |
| Rorc | 2557.825223 | 0.751721127 | 0.118332 | 6.352648932 | 2.12E-10 | 3.27156E-08 |
| Blk | 3075.942673 | 0.834934771 | 0.119593 | 6.981490603 | 2.92E-12 | 7.08357E-10 |
| Ccr6 | 470.5011763 | 1.064628255 | 0.145442 | 7.319938762 | 2.48E-13 | 9.69389E-11 |
| Scart2 | 607.7134197 | 1.167265425 | 0.205535 | 5.679161575 | 1.35E-08 | 1.22672E-06 |
| Arnt2 | 480.0998444 | 1.295461383 | 0.189359 | 6.841288483 | 7.85E-12 | 1.67278E-09 |

